# Exact-match search with functional variant prediction enables automated DNA screening

**DOI:** 10.1101/2024.03.20.585782

**Authors:** Dana Gretton, Brian Wang, Rey Edison, Leonard Foner, Jens Berlips, Theia Vogel, Martin Kysel, Walther Chen, Francesca Sage-Ling, Lynn Van Hauwe, Stephen Wooster, Helena Cozzarini, Benjamin Weinstein-Raun, Erika A. DeBenedictis, Andrew B. Liu, Emma Chory, Hongrui Cui, Xiang Li, Jiangbin Dong, Andres Fabrega, Christianne Dennison, Otilia Don, Cassandra Tong Ye, Kaveri Uberoy, Ronald L. Rivest, Mingyu Gao, Yu Yu, Carsten Baum, Ivan Damgard, Andrew C. Yao, Kevin M. Esvelt

**Author notes:** For correspondence: Kevin M. Esvelt, PhD, Massachusetts Institute of Technology, Building E15-354, 77 Massachusetts Ave, Cambridge, MA 02139 USA. equal contribution (Dana Gretton, Brian Wang).

## Abstract

Custom DNA synthesis underpins modern biology, but controlled genes in the wrong hands could threaten many lives and public trust in science. In .1992, a virology-trained mass murderer tried and failed to obtain physical samples of Ebola; today, viruses can be assembled from synthetic DNA fragments. Screening orders for controlled sequences is unreliable and expensive because current similarity search algorithms yield false alarms requiring expert human review. Exact-match search can achieve perfect specificity among short known sequences by detecting subsequences unique to controlled genes, but can be trivially evaded by incorporating mutations. Here we rescue exact-match search by additionally screening for predicted functional variants of pseudo-randomly chosen subsequences that aren’t found in known unregulated genes. To experimentally assess robustness, we protected nine windows from the M13 bacteriophage virus, then invited a “red team” to launch up to 21,000 attacks at each window and measure the fitness of their designed mutants. We identified defensible windows from regulated pathogens, built a test database that our experiments indicate will block 99.999% of functional attacks, and verified its sensitivity against redesigned enzymes. Exact-match search with functional variant prediction offers a promising way to safeguard biotechnology by automating DNA synthesis screening.

**Summary:** Searching for exact matches to pre-computed functional variants unique to controlled genes enables sensitive, secure, and automated DNA synthesis screening.

## Introduction

The COVID-19 pandemic demonstrated that society is profoundly vulnerable to new transmissible biological agents, even as inexpensive DNA synthesis, assembly methods, and detailed reverse genetics protocols have made harmful viruses accessible to a large and growing number of technically skilled individuals^1–4^. Recent publications strongly suggest that future advances will provide genomic blueprints and reverse genetics protocols for credible pandemic agents^4–15^.

Fortunately, most individuals skilled enough to assemble viruses with reverse genetics must obtain synthetic DNA from a commercial supplier (Fig. 1a). Members of the International Gene Synthesis Consortium (IGSC), an industry group, are committed to screening DNA synthesis orders above a certain length^16^.

**Figure 1.**
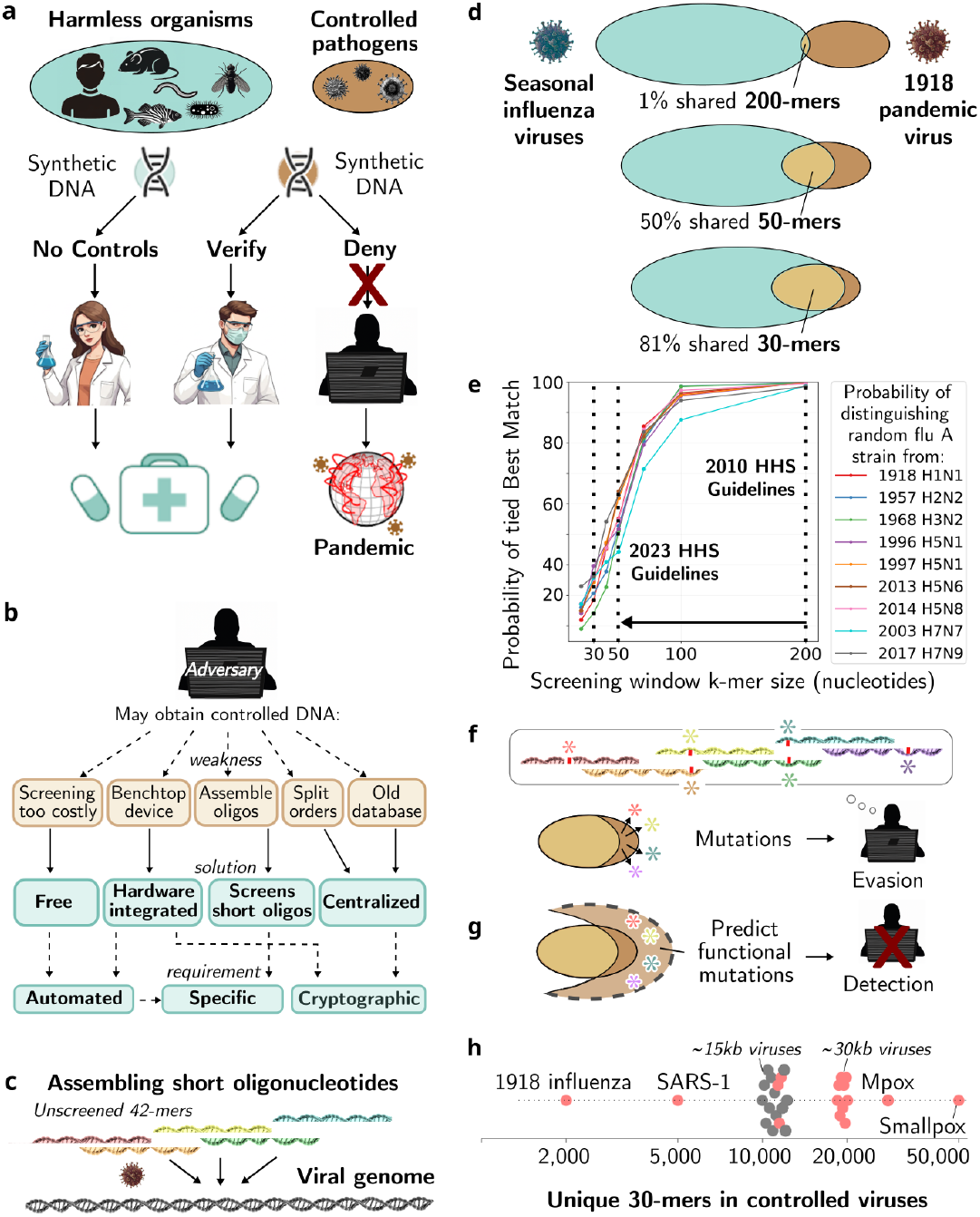
Achieving robust DNA synthesis screening. **a)** The goal of screening is to allow anyone to obtain nucleic acids from harmless organisms, but require verification and reporting before shipping oligonucleotides matching controlled pathogens in order to prevent misuse. **b)** Adversaries can evade current screening by exploiting weaknesses. Not all providers screen due to the cost, benchtop devices cannot wait for human review, unscreened short oligos can be assembled, and decentralized sources are vulnerable to split orders and failures to update. Centralized screening can only preserve the privacy of customer orders with cryptography, which will require a highly specific and efficient search algorithm. **c)** Short unscreened oligonucleotides can be used to generate controlled sequences such as viral genomes. **d)** Screening for subsequences unique to controlled agents (brown) achieves perfect specificity among known sequences. As *k*-mer length decreases, there are fewer unique sequences within any given controlled agent, especially for controlled agents such as 1918 influenza that are closely related to many unregulated agents, but there are still enough unique sequences for exact-match search to detect. **e)** BLAST-based screening is worse than

The IGSC deserves praise for voluntarily prioritizing safety because screening is becoming costlier as the price of synthetic DNA falls^17,18^. Unfortunately, more than two thirds of gene synthesis firms are non-members. A modern equivalent of 1992’s virology-trained terrorist^19^ can plausibly obtain potential pandemic viruses by ordering the DNA from a company not listed on the IGSC website.

Even if all providers did screen requests, adversaries could obtain DNA sufficient to generate a pandemic virus by assembling oligonucleotides shorter than the minimum length for screening; by ordering non-overlapping pieces from multiple suppliers (a split-order attack); or by swiftly placing orders for newly identified pandemic viruses to exploit databases that are slow to update, among other tactics (Fig. 1b). Recent red-teaming confirmed many of these weaknesses, obtaining fragments collectively sufficient to assemble infectious 1918 influenza virus, without any authorization requests, from nearly every firm tested^20^. Benchtop machines enabling on-site synthesis create another vulnerability^17,21,22^: not only must each device be updated whenever a new threat is identified, but the screening system cannot be stored locally on the device lest it be interrogated and used to build screening-evasion software allowing others to obtain controlled sequences without being detected. Verifiably screening all commercial and benchtop DNA synthesis^23^ demands an automated, centralized, and privacy-preserving approach that reliably detects controlled sequences. Critically, such a system must trigger negligibly few false alarms to avoid time-consuming human review of each flagged sequence.

Current BLAST-based screening, which determines the known sequences that represent the Best Matches to each subsequence window of an order, is too nonspecific to be automated. While relatively accurate for 200-mer windows, screening 50-mer and 30-mer windows to prevent oligonucleotide assembly (Fig. 1c) or mutation correction dramatically degrades Best Match specificity: whereas nearly all 200-mers in the 1918 pandemic influenza virus are unique to that strain, fully 50% of 50-mers and 81% of 30-mers are shared with unregulated strains (Fig. 1d). As a result, recent U.S. guidelines specifying 50-mer windows^24^ cause many Best Match false alarms; most unregulated influenza A strains will trigger at least one such alarm (Fig. 1e), necessitating costly expert review.

In contrast, exact-match search can exclusively screen for short k-mers unique to controlled agents, achieving perfect specificity among all known sequences, even for controlled agents closely related to unregulated organisms (Fig. 1d). This specificity is impossible with BLAST-based approaches, which necessarily generate numerous false alarms for shorter sequences (Fig. 1e). However, exact-match screening can be readily evaded by incorporating functional mutations into each k-mer (Fig. 1f).

We hypothesized that exact-match search could be rescued by predicting functional variants of pseudo-randomly chosen subsequence windows to detect mutated or redesigned sequences, then curating the resulting database by removing any predicted variant that matches a known unregulated sequence (Fig. 1g). Instead of finding the Best Match to every window in an order, we can check for matches to unique windows and predicted variants. Virtually all controlled agents harbor thousands of unique 30-mers, making them readily defensible (Fig. 1h, Extended Data Fig. 1). Here we analyze the specificity that can be achieved with exact-match search and experimentally assess its adversarial robustness when supplemented with basic functional variant prediction; our companion paper analyzes real-world exact-match search at assessing shorter *k*-mer windows because it must flag any window for which the Best Match includes a controlled sequence, even one identical to an unregulated sequence. Since few controlled and unregulated influenza A strains share 200 base pair windows, but most strains share at least one 50 base pair window with a controlled strain, they will yield many Best Match false alarms under the updated 2023 HHS guidance recommending 50-mer screening. **f)** The primary weakness of exact-match screening is that it detects only wild-type sequences, allowing even a few mutations to render controlled sequences undetectable. **g)** Screening for predicted functional variants not found in unregulated sequences may restore robustness while preserving specificity. **h)** Uniqueness of 30-mers found in U.S. select agent (red) and other controlled (grey) viruses. specificity and the system-level performance of an operational privacy-preserving cryptographic implementation (Fig. 2a).

**Figure 2.**
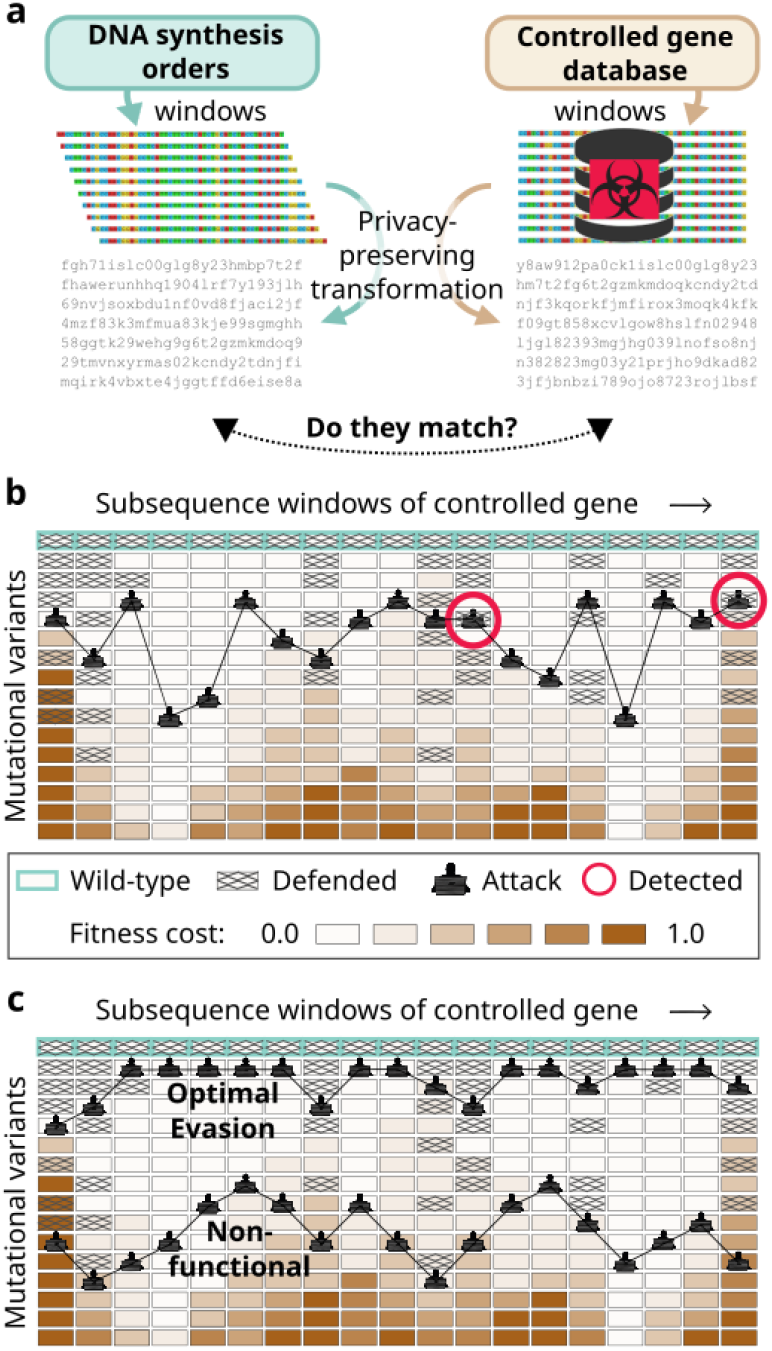
Rescuing exact-match search with predicted functional variants. **a)**Screening relies on detecting matches between subsequences from DNA synthesis orders and from controlled sequences. Efficient exact-match search permits the use of cryptography to preserve the privacy of both orders and the database^25^. **b)** To evade detection by strategically choosing functional variants, the adversary must choose a mutated subsequence for every window across the coding sequence of the controlled agent without picking one in the database, as shown. This is maximally challenging when the database is populated with predicted functional variants of windows that tolerate few mutations. Detection becomes more probable as the defender’s predictive capacity increases relative to the attacker’s. **c)** An attacker who knows which windows are defended can evade imperfect screening. Keeping the database remote and choosing windows and variants quasi-randomly forces adversaries to guess, likely including so many mutations that the resulting hazard is no longer functional.

## Results

Suppose that an adversary seeks to obtain a regulated agent W by incorporating mutations (Fig. 1f). For each mutated subsequence window *w*_i_, there are three possible outcomes:

1. *w*_i_ is present in the controlled gene database, and the synthesis order is rejected and logged
2. *w*_i_ escapes detection, but imposes a fitness cost *c*_i_ that reduces functionality
3. *w*_i_ escapes detection at negligible fitness cost

Success requires the adversary to achieve the third outcome for most *w*_i_ to preserve function. We define the *random adversarial threshold R* as the probability that an adversary with perfect knowledge of the fitness of each variant—but ignorant of which windows and variants are defended—will be detected upon attempting to synthesize functional W.

In theory, defending all variants that do not completely abolish the function of W at one essential window *w*_i_ can perfectly thwart the adversary, achieving *R*=1. In practice, fitness prediction is imperfect, but *R* can still be maximized by defending many of the windows predicted to be least tolerant of mutations and optionally adding new windows when an attempt is detected (Fig. 2b, Extended Data Fig. 2).

An adversary with superior predictive capacity who learns which function-prediction algorithms are used can evade screening by choosing the highest-fitness undefended variant known to them for each window (Fig. 2c). By choosing which windows and variants to defend quasi-randomly, we can force the adversary to heavily mutate all windows throughout the hazard in order to evade detection, greatly reducing their odds of obtaining functional W (Extended Data Fig. 2).

### Choosing a window size while maintaining specificity

A key benefit of exact-match screening is the ability to curate the database by removing all peptides and k-mers that match unregulated sequences from sequence repositories, thereby perfectly eliminating known false positives (Fig. 3, Extended Data Fig. 2). We distinguish such sequences from controlled sequences and close relatives using taxonomic classification, keywords, and counting the number of windows that match the hazard, among others. As our companion paper demonstrates using customer data, exact-match predictive screening using a large database will seldom if ever flag known unregulated sequences^2^.

**Figure 3.**
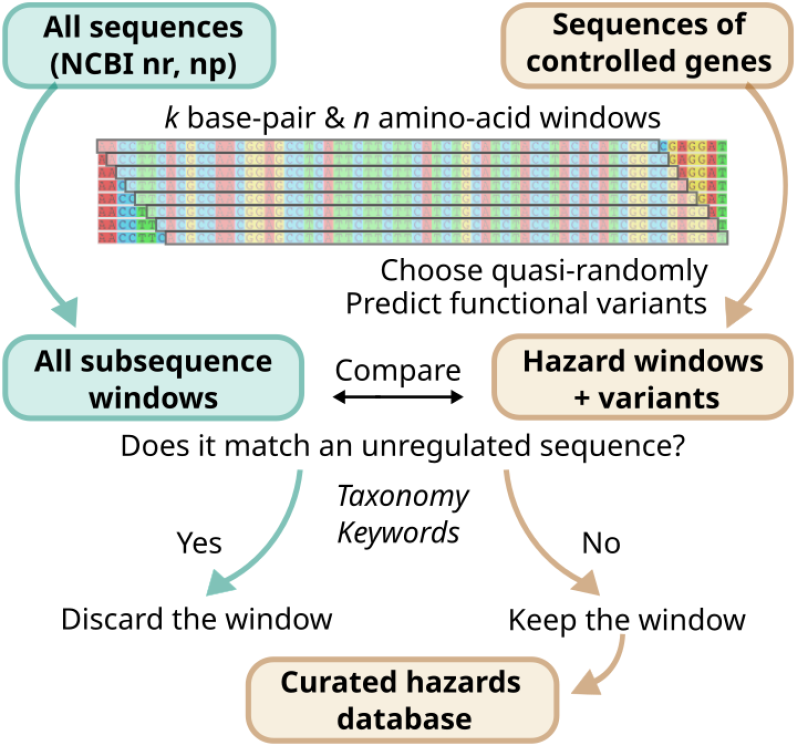
Building the database of sequences unique to controlled agents. Subsequences from controlled sequences and predicted functional variants are compared to similarly-sized windows from NCBI repositories. Matches to repository sequences that are unregulated—as determined by taxonomy, keywords, and fraction of matching windows—are discarded to reduce the false alarm rate.

However, synthetic sequences absent from natural organisms—which are commonly employed in deep mutational scanning and directed evolution experiments as well as designed proteins—will still randomly trigger false alarms, as will natural sequences not deposited in public databases. Random false alarms will occur at a frequency determined by the total amount of novel DNA synthesized, the number of sequences in the database, and the window length (SI Appendix A).

Historical efficiency improvements^26^ and market projections suggest that global annual synthesis demand may rise to as much as 10^15^ base pairs in a decade. While almost certainly an overestimate, this is counterbalanced by the fact that functional biopolymer sequences are *not* randomly distributed^27^: peptide frequencies are highly biased by amino acid composition and functional constraints^28–30^. For our initial experiments aimed at measuring *R*, we chose to screen peptides of length 19 because searching for 10^7^ functional variants for each of 1,000 regulated agents would yield one truly random false alarm per 10^15^ base pairs of DNA.

### Experimentally testing sensitivity

To test the efficacy of exact-match predictive screening against deliberately introduced mutations and learn which types of windows are most easily defended, we selected the harmless M13 virus that infects *E. coli* as a “controlled” genome. To investigate which peptide windows might be most relevant to defense, we analyzed all length 19 peptide subsequences using fuNTRp, a computational tool that categorizes residues within proteins as “neutral” if they likely tolerate most mutations, “rheostat” if they suffer reduced fitness from many but not all mutations, or “toggle” if nearly any mutation is deleterious.^31^ Lower neutral scores indicate less mutational tolerance with preserved functionality, suggesting enhanced utility as defensive screening targets. From four required M13 proteins (Extended Data Fig. 3), we selected nine total windows with fairly low to very low neutral values and a range of rheostat and toggle scores (Extended Data Fig. 4, Extended Data Table 1).

Next, two “blue team” members constructed databases of 10^3^ to 10^7^ predicted functional variants for each window using a Metropolis-Hastings algorithm^32^ that combined fuNTRp scores with the BLOSUM62 amino acid substitution matrix. While more sophisticated variant-effect predictors exist, they are typically optimized for predicting effects of single mutations in human genes^33^, not multiple mutations in pathogenic proteins. We chose this minimal fuNTRp+BLOSUM62 approach in part to test whether the inherent asymmetry in the screening problem—where attackers must evade screening at all windows while defenders need only succeed in select cases—provides sufficient leverage for a less sophisticated predictive model to prevail in the worst-case scenario when attackers have access to superior prediction tools^34,35^ (SI Appendix B).

“Red team” members experimentally tested the security of exact-match predictive search by designing and launching up to 21,000 attacks at each of the nine windows using combinatorial rational design (Fig. 4a). They chose to order oligonucleotide pools with all possible combinations of the four most common substitutions at the six positions with the highest neutral scores, pairwise substitutions of all amino acids at those six positions, and all possible single substitutions. They generated libraries of variants by molecular cloning and measured the effects of each variant on phagemid replicative fitness via complementation.

**Figure 4.**
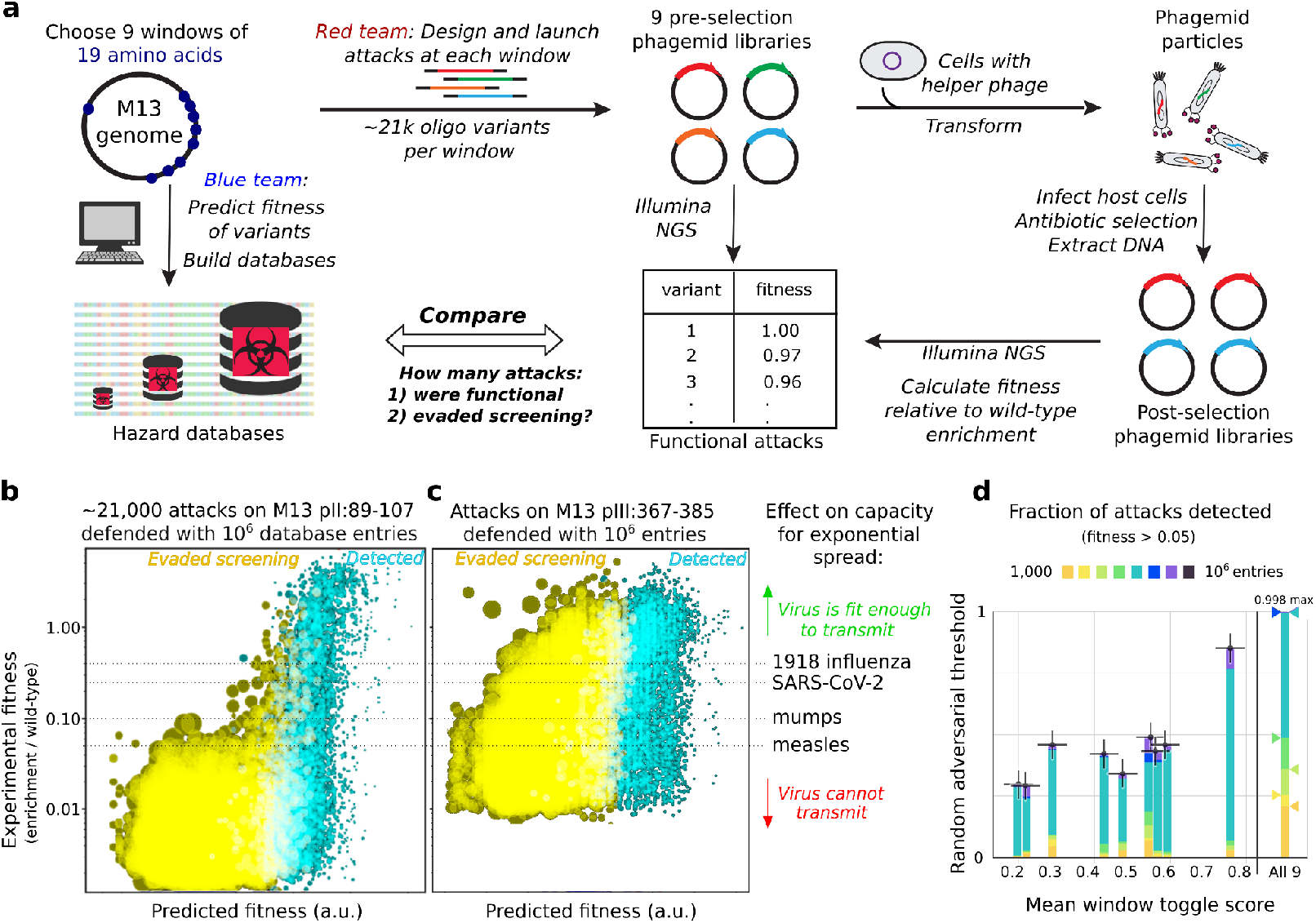
Incorporating mutations into the genomic blueprint of a virus cannot readily escape screening. **a)** Team members built defensive databases by predicting functional variants for nine different windows in the genome of M13 bacteriophage. Others launched∼21,000 attacks at each window by synthesizing variants with up to six amino acid changes and using a phagemid assay to measure the fitness of each variant, which we defined as enrichment relative to the wild-type sequence. **b)** At the most defensible window, located within the M13 pII endonuclease, 92% of attacks yielding variants with fitness 0.06 and above were thwarted by screening. Smaller dots correspond to sequences with fewer mutations. **c)** At a moderately defensible window located within the M13 pIII receptor-binding protein, 49% of such attacks were thwarted, underscoring the importance of window choice. Potential pandemic pathogens can tolerate only so many mutations impairing fitness before they are no longer capable of sustained transmission. The corresponding fitness lines depict these threshold values for 1918 influenza (R_0_~2.5), SARS-CoV-2 (R_0_~4), mumps (R_0_~10), and measles (R_0_~18), which is the most contagious virus known. **d)** The fraction of attacks detected, which corresponds to the random adversarial threshold, as a function of the average fuNTRp toggle score for each of the nine windows for various database sizes (1000, 2000, 5000, 10^4^, 5×10^4^, 10^5^, 5×10^5^, 10^6^) using a fitness cutoff of 0.05 (sufficient to prevent the sustained spread of measles). Combinatorial screening of all nine windows (right) detected virtually 100% of individually functional attacks.

We defined “functional” variants as those with a measured fitness of at least 0.05 relative to wild-type, which is the level at which the most infectious virus known can no longer spread in an unprotected population^36^. Since approximately 50% of variants chosen by the attackers met this standard, the red team effectively launched∼10^33^ combinatorial, individually functional attacks on the nine windows in the database. Of the attacks on the most defensible window, 85.2% were detected by a database with 10^6^ entries (Fig. 4b). That is, even an adversary with perfect knowledge of subsequence fitness who already possesses the other 99% of the wild-type M13 genome sequence was likely to be detected and thwarted at just this one window. While other windows were less defensible, most still blocked∼30-50% of attacks (Fig. 4c). When the nine M13 windows were defended by 100,000, 1 million, or 10 million variants, 99.3%, 99.7%, and 99.8% of individually functional attacks were detected at one or more windows (Fig. 4d). These results underscore the extreme difficulty of obtaining a functional hazard by incorporating mutations to evade exact-match predictive search at all relevant windows.

Moreover, windows do not exist in isolation: an attack featuring mildly deleterious genomic mutations in two different windows that separately reduce fitness to 0.20 typically has a *combined* fitness of 0.04 (or less)^37,38^, and consequently will not be functional (Extended Data Fig. 5). We simulated combinatorial attacks by randomly combining functional variants at each window into 10^10^ M13 phage genomes, multiplying the fitness values for all nine windows, and discarding those with a combined fitness below 0.05, forcing variants to maintain higher individual fitness at each window to produce viable genomes. Because high-fitness variants were more readily predicted by the defenders (Extended Data Fig. 6), defending just 50,000 variants per window successfully detected 99.94% of combinatorially functional attacks by the red team.

Most importantly, defending 20 windows equivalent to the geometric mean of the nine from M13 with just 50,000 database entries per hazard would block 99.999% of attacks seeking to obtain a pandemic virus as contagious as anything known to science (Fig. 5a, Extended Data Fig. 7). In practice, the database will typically include 100 or more windows for each controlled agent.

**Figure 5.**
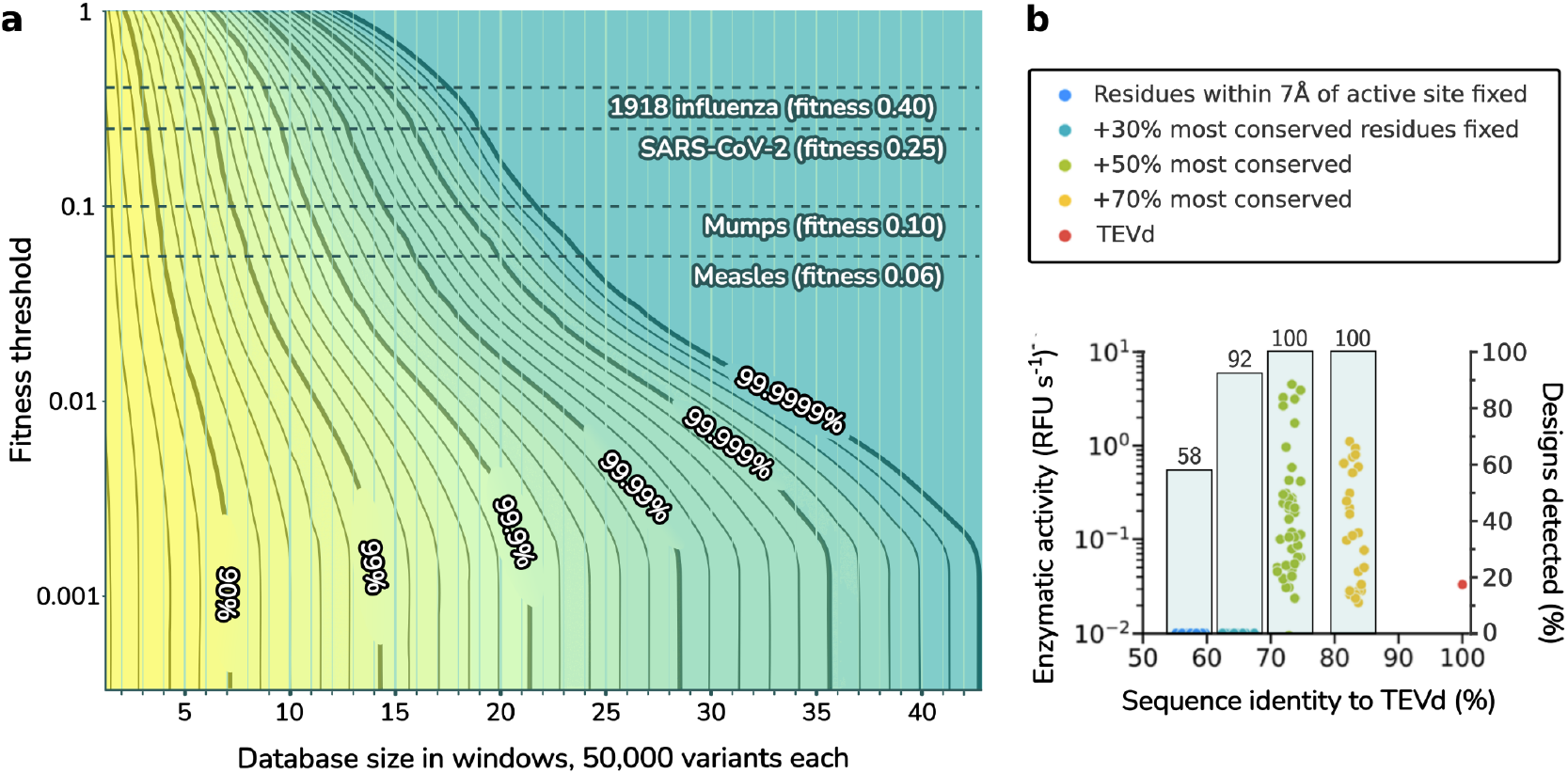
Adversarial and machine-learning-generated designs are reliably detected. **a)** The cumulative effect of safeguarding more or fewer windows can be extrapolated using powers of the geometric mean of escapee curves (Extended Data Figs. 6-7). The Random Adversarial Threshold is plotted as a function of both simulated number of windows protected with 50,000 variants, and the fitness at which the hazard is no longer considered functional. Contours show lines of equal protection as trade-offs between the minimum fitness tolerated and number of windows protected. As an illustration, 16 mean windows would block 99.99% of attempts to synthesize and spread a virus as contagious as measles, or 99.999% of attempts to make a viable pathogen as infectious as SARS-CoV-2. In practice, the database includes 100 or more windows for each virus. **b)** Detection of redesigned functional and nonfunctional TEVd protease variants generated with ProteinMPNN by crude BLOSUM62 prediction, grouped by identity to the wild-type enzyme^39^. All functional redesigns were detected despite the use of inferior variant prediction tools. Future databases will incorporate ProteinMPNN and improved tools for prediction.

Since the red team knew exactly which windows they needed to mutate, and was able to test functional variants without needing to worry about being detected at any other windows, these results strongly suggest that the random adversarial threshold *R* can approach 1.0 unless the attacker has notably superior predictive capability (SI Appendix B).

Window selection and database allocation can be optimized based on estimated defensibility. Since adversaries can obtain sequence variants appearing in related viruses that are not regulated, defenders should cross-reference genomes and prioritize some windows lacking functional equivalents in relatives. For optimal allocation of limited database entries, the fuNTRp “toggle” score, or predicted sensitivity to mutation at each position^31^, appears most predictive of *R* across the nine windows tested (Extended Data Fig. 8). Analysis of how toggle scores relate to the trade-off between false positive and false negative rates suggests that such proxies for defensibility can inform the optimal number of database entries needed for each window (Extended Data Fig. 9).

### Sensitivity against adversarial attacks

Having established the robustness of exact-match predictive search against evasive strategies focused on introducing mutations, we sought to determine effectiveness against a variety of attacks that defeat both alignment and exact-match search. We built and curated a controlled sequence database from the U.S. Select Agents and Australia Group pathogens (Methods). To prevent adversaries from ordering and assembling short oligonucleotides, we additionally included all 30-mer DNAs as well as 42-mers with all single mutations.

To test sensitivity against attempted evasion, we wrote scripts that used several distinct strategies to design gene synthesis orders for sequences from 1918 influenza and ricin, both U.S. select agents, that reliably evade detection by both BLAST and straightforward exact-match screening for 50-mer nucleotides and 16-amino-acid peptides, yet can be assembled into functional hazards in no more than two laboratory steps using standard protocols^20^. Testing revealed that our implementation of exact-match predictive search detected each controlled sequence disguised using these scripts without flagging unregulated sequences.

### Quantifying sensitivity-specificity tradeoffs through bioinformatic analysis

Exact-match predictive screening can trade-off between specificity and sensitivity in a controlled manner. At maximum specificity, only k-mers unique to controlled pathogens are flagged, while sequences shared with related but unregulated pathogens are ignored. An analysis of 30-mers from 49 controlled viruses against a comprehensive dataset of unregulated viral 30-mers revealed that the majority of viruses maintained nearly 100% 30-mer uniqueness, with only SARS, 1918 influenza, variola, and mpox exhibiting less than 50% unique sequences (Extended Data Fig. 1). Notably, all controlled viruses except SARS and 1918 influenza possessed more than 9,000 unique 30-mers, with these exceptions showing high sequence similarity to other coronaviruses and influenza viruses, respectively.

To enable detection of controlled pathogens with few unique sequences, screening for additional k-mers shared with a handful of unregulated relatives can increase sensitivity at the expense of specificity. For example, an adversary could obtain a pandemic influenza virus by first ordering all sequences shared with unregulated pathogens, then seeking to acquire the relatively tiny number of sequences unique to the pandemic strain. The first step could be made much more difficult by additionally screening for and denying DNA orders shared by the pandemic virus and some rare unregulated influenza strains.

In the companion paper, we report the results of specificity testing on real-world DNA synthesis orders, which identified zero false alarms from orders corresponding to known sequences in public databases^40^. Just 1 in 5,000 orders corresponded to a novel strain of a related virus not yet added to public databases. These findings demonstrate that exact-match screening can maintain high sensitivity even at maximum specificity.

### Robustness against protein design tools

In principle, machine learning tools may be capable of altering enough residues in controlled proteins to evade screening while preserving function. To challenge exact-match predictive search against enzyme redesign, we built a controlled sequence database to defend TEV protease, which was recently redesigned with ProteinMPNN^39^, and screened all reported designs. Fully 100% of designs with nonzero activity were detected, despite the comparative crudity of fuNTRp+BLOSUM62 prediction (Fig. 5b). Incorporating more advanced design tools into the database generation pipeline will allow screening to detect redesigned threats even more reliably (SI Appendix B).

## Discussion

For decades, obtaining dangerous pathogens required access to specialized facilities or infected samples. Today, any suitably skilled individual with a credit card can legally order and assemble the building blocks of life to generate toxins and viruses. The technical limits of screening via homology search have precluded widespread adoption and policy mandates, leaving the door ajar for misuse—only prevented by the temporary dearth of known pandemic-capable viruses accessible via synthetic DNA. Unlike BLAST-based approaches that generate numerous false alarms requiring expert review, screening for exact matches to subsequences unique to controlled agents, combined with functional variants, eliminates false positives from known sequences while retaining high sensitivity. The success of Kraken^41^ and related tools demonstrates that exact-match search can effectively safeguard biotechnology when properly implemented. This crucial advance enables fully automated screening—fast, reliable, and achievable in benchtop devices where human review is impossible.

While our experimental tests demonstrate robustness against adversarial attacks, increasingly sophisticated protein design tools will allow greater sequence divergence while preserving function, even at conserved windows^39,42^. To stay ahead, we will need to incorporate the same machine learning approaches that enable more effective protein design into our variant prediction algorithms to preemptively include these newly accessible sequences in our database.

There are theoretical limits: *de novo* design tools may eventually generate so many functional variants of proteins that no database could feasibly contain them all. Someday, orders for designed toxins—particularly if split among multiple providers to evade structure- and function-based prediction—will likely evade detection by all screening systems. Thankfully, the far greater danger posed by pandemic-capable viruses should remain at bay for many years due to the complexity of redesigning an entire self-replicating pathogen without extensive laboratory validation.

The most urgent need to implement universal synthesis screening may arise from the application of specialized design tools to viral entry proteins, which will eventually predict which variants can evade current population immunity while retaining their function. These tools are being developed with the best of intentions, but if made publicly credible and accessible, they will unavoidably enable malefactors to equip viral backbones with thousands of distinct entry proteins—generating many novel pandemic viruses, each of which requires its own targeted vaccine. Because synthetic DNA is far more accessible than isolating pathogens from natural sources, controlling access to viral backbone sequences will be increasingly important.

Towards this goal, our companion paper describes a free, automated cryptographic screening and permissions system that implements this approach while preserving the privacy of both orders and database entries. By reliably and automatically allowing only non-controlled sequences to proceed unimpeded, while consistently requiring proper biosafety approvals for controlled material, this system will naturally establish a new norm where researchers routinely include appropriate certificates when ordering potentially regulated DNA. Given revised legal frameworks that mandate screening, perhaps by regulating DNA fragments encoding select agent viruses as select agents themselves, this system could become universal, defending us against the short-sequence and split-order attacks that so readily obtained all nucleic acids required to generate the 1918 influenza virus. As we learn to write in the language of life, we must keep the tools for banishing disease from becoming instruments for synthesizing plagues.

## Methods

### Bioinformatic analysis

All influenza genomes were downloaded as .fasta files from NCBI Virus and filtered to remove any with fewer than 30 nucleotides. Segments of controlled or pandemic genomes were downloaded and combined into single .fasta files. The 1918 genome comprised A/Brevig Mission/1/1918(H1N1) except segment 4, from A/South Carolina/1/1918(H1N1).

### Window selection: M13 peptides

The proteins of bacteriophage M13 (Accession NC_003287.2) were analyzed using fuNTRp^31^, which scores each amino acid position by the likelihood that it will accept many (neutral), some (rheostat), or few (toggle) different substitutions without disrupting protein function. We used fuNTRp to identify peptide windows with few predicted neutral positions and varying numbers of toggle and rheostat positions across and within proteins (Extended Data Table 1).

### Blue team: Defending the nine windows of M13

All single-nucleotide mutations were included for reasons of caution. For the remaining entries, a Metropolis-Hastings algorithm was used to select combinations of mutations predicted to minimally impair fitness. The mutation tolerance scores estimated by fuNTRp were combined with the BLOSUM62 matrix to generate a probable cost for every possible substitution, with costs multiplied for multiple-mutation combinations.

*Red team: Procedurally generating variants for each 19-aa peptide window*

1. We included the wild-type sequence (1)
2. We included all one-mutants at each position (19×19 = 361)
3. At the six positions predicted to be most neutral, we added all combinations of the four predicted least pathological substitutions according to BLOSUM62 (5^6^ = 15625) (overlaps with one-muts and WT at 4×6+1=25)
4. As negative controls (not attacks), we included up-to-six mutants of neutral positions using the two most pathological substitutions according to BLOSUM62 (3^6^ = 729) (overlaps with one-muts and WT at 2×6+1=13)
5. We added all pairwise combinations of all possible substitutions at the six most neutral positions (19^2^×15 pairwise combinations = 5415) (overlaps with 4 most tolerated at 4^2^×15=240) (overlaps with 2 most pathological at 2^2^×15=60) (included in case the defenders did not block all of them)

Total: 1 + 361 - 13 + 15625 - 25 + 5415 - 240 - 60 = 21,793 peptide variants at each window.

### Construction of phagemid libraries

Oligo libraries comprising variants for each 19aa peptide window were synthesized as a pool by Twist Bioscience. Individual libraries were amplified by PCR and ligated into a phagemid backbone—encoding an ampicillin resistance gene, containing an M13 phage origin of replication, and designed for library variant expression upon induction by IPTG—using NEBuilder Hifi DNA Assembly Master Mix (NEB, E2621L). All libraries were then precipitated with isopropanol, transformed into electrocompetent DH5α cells (NEB, C2989K), and plated on 2XYT-carbenicillin-1% glucose; after overnight growth at 37 °C, colonies were counted to ensure >50-fold library coverage. Colonies were scraped with 2XYT and plasmid DNA extracted with the ZymoPURE II Plasmid Maxiprep Kit (Zymo Research, D4203); the extracted plasmid DNA was then precipitated with isopropanol. These plasmid libraries constitute the “pre-selection libraries.”

### Construction of helper cells

M13cp^43^, a plasmid containing all M13 phage genes but with a p15a origin and a chloramphenicol resistance gene replacing the phage origin of replication, was used to construct helper plasmids. Primer pairs were designed for the precise deletion of genes I, II, III, and IV from M13cp following PCR amplification and ligation using the In-Fusion Snap Assembly Master Mix (Takara Bio, 638944). The resulting helper plasmids were transformed into DH5α competent cells (NEB, C2987H), yielding four individual helper cell lines (M13cp-dg1, M13cp-dg2, M13cp-dg3, and M13cp-dg4). The helper cells were made electrocompetent for subsequent same-day transformations. Helper cells are capable of extruding phagemid particles when transformed with a phagemid library variant with a functional gene (complementing the missing phage gene in the helper plasmid) and origin of replication (Extended Data Figure 3). DNA sequences of helper plasmids and phagemids expressing wild-type proteins are available on Addgene.

### Phagemid growth

Phagemid libraries were transformed into their corresponding helper cells (nucleic acid variant libraries were transformed into M13cp-dg3) by electroporation and plated on 2XYT-carbenicillin-chloramphenicol-1% glucose. After overnight growth at 37 °C, colonies were counted to ensure >15-fold library coverage. Colonies were scraped with 50 mL 2XYT, the bacterial pellet washed sequentially 3x with 50 mL 2XYT, then a 1:1000 dilution used to inoculate a 50 mL phagemid growth culture in 2XYT with maintenance antibiotics and 1% glucose. The culture was grown to OD_600_ = 0.5 with shaking at 37 °C and 250 rpm, at which point the culture was centrifuged and the media replaced with 2XYT containing maintenance antibiotics and 1 mM IPTG. The culture was grown for 16 h at 37 °C and 250 rpm, after which phagemid-containing supernatants were collected by culture centrifugation and filtration through a 0.22 μm filter.

### Phagemid infection

Phagemid-containing supernatants were added to 2.5 mL S2060 cells (streptomycin-resistant, Addgene #105064) grown to OD_600_ = 0.5 and allowed to infect at 37 °C and 250 rpm for 1 h. The resulting infected cultures were plated on 2XYT-carbenicillin-streptomycin-1% glucose to select for phagemid-containing cells. After overnight growth at 37 °C, colonies were scraped with 50 mL 2XYT and plasmid DNA extracted with the ZymoPURE II Plasmid Maxiprep Kit (Zymo Research, D4203). These plasmid libraries constitute the “post-selection libraries.”

### Illumina NGS sequencing

Pre- and post-selection libraries were prepared for illumina NGS sequencing by sequential PCR amplification. PCR amplification was first performed with PrimeSTAR GXL Premix (Takara Bio, R051A) to attach Nextera-style adapter sequences, followed by a second PCR amplification to attach library-specific barcodes and the p5 and p7 indices. Following PCR purification, library concentrations were quantified with qPCR using the NEBNext Library Quant Kit for Illumina (NEB, E7630S), and pre- and post-selection libraries were combined as two pools. Libraries were pooled such that libraries were present in equimolar quantities corrected for library size. Libraries were submitted to the MIT BioMicro Center for MiSeq Illumina sequencing (v3, 2 x 300 bp paired-end).

### Controlled sequence database generation

We began by generating a curated database comprising subsequence windows from U.S. Select Agents, Australia Group hazards, and functional variants not found in unregulated genes and genomes (Fig. 2a). We included all 19-amino-acid peptides and all 42-mers as well as 30-mers from each hazard; reliably assembling hazards from smaller DNA pieces is much more challenging^44,45^. To detect adversarial attempts to obtain functionally equivalent mutants of known hazards, we quasi-randomly selected windows to defend and used a variant effect predictor to compute and add millions of variants per window^31,34,46–50^.

Next, we curated database entries to remove peptides and k-mers matching unregulated sequences found in GenBank nr/nt and protein nr using taxonomic classification, keywords, and exact match quantification, as well as entries with a low Shannon entropy (Fig. 3). Curation perfectly avoids flagging unregulated sequences in public databases, eliminating all known, nonrandom false alarms. As detailed in our companion paper, the efficiency of exact-match search permits the use of cryptography to protect the privacy of DNA synthesis orders sent to be screened while shielding the database from interrogation^25,51^.

### Bioinformatics specificity analysis

Representative whole-genome sequences from pathogenic strains of selected human and animal viruses were obtained from the NCBI nucleotide database. For each viral sequence, we generated all possible 30-mers. In parallel, we downloaded the complete U-RVDB viral sequence database (v29.0) and generated all possible 30-mers, excluding sequences from controlled viruses based on a curated list of NCBI taxonomy IDs. To identify virus-unique sequences, we computed the set difference of each controlled-virus 30-mer set against the unregulated viral 30-mer set.

## Acknowledgements

We thank S. Golas for assistance with scripting and AWS, K. Sumida for providing data on ProteinMPNN designs, and E. Soice for figure design support.

## Author contributions

K.M.E., D.G., and A.C.Y. conceived the study; L.F., R.R., M.G., Y.Y., C.B., I.D., and A.C.Y. performed the security assessments, D.G. wrote software to implement the search method to defend M13; E.A.D. designed the attacks against M13; B.W. designed and performed all laboratory experiments involving M13; E.C. analyzed sequencing data; D.G. performed the sensitivity analyses; D.G. wrote the prototype database software with assistance from C.D., O.D., T.Y., K.U., and A.F.; the initial screening prototype was developed by H.C., X.L., J.D., M.G., and Y.Y.; professional software development was overseen by L.F. and J.B. and performed by L.F., J.B., T.V., M.K., W.C., F.S-L., L.V.H., S.W., and B.W-R.; and the attacks that defeat alignment and conventional exact-match search were devised and tested by R.E. and K.M.E. All authors contributed to writing the paper.

## Funding

We are grateful for financial support from the Open Philanthropy Project (to MIT, Aarhus University, and SecureBio), an anonymous philanthropist from mainland China (to Tsinghua University), the Aphorism Foundation (to MIT and SecureBio), and Effective Giving (to MIT and SecureBio). The funders had no role in study design, data collection, data analysis, data interpretation, or writing of the report. To preserve international neutrality, no government funds were used to support this project.

## Competing interests

K.M.E. and D.G. are authors of PCT/US2021/014814 filed by the Massachusetts Institute of Technology. All authors share an interest in preventing future pandemics.

## Data availability

Raw experimental data and analyses of the fitness of mutant M13 phages and database entries used for defense are available via Figshare.

## Code availability

Code is available at https://github.com/dgretton/RAT-DNA-screening. fuNTRp^31^ is available on request from Maximilian Miller or at https://bromberglab.org/project/funtrp/.

Supplementary Information is available for this paper.

Correspondence and requests for materials should be addressed to.

**Extended Data Fig. 1.**
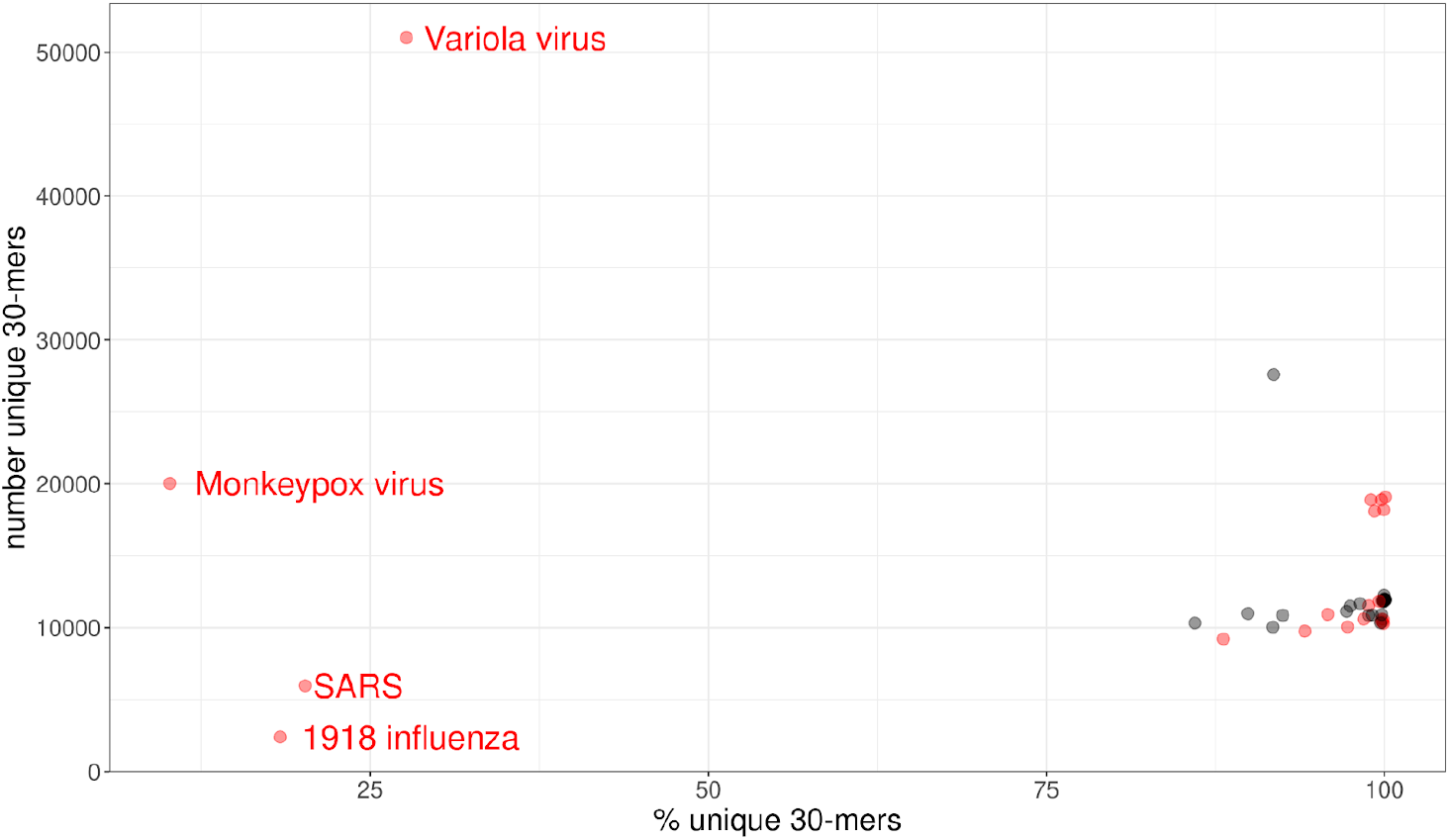
Identification of virus-unique 30-mers. SecureDNA screens DNA synthesis orders by identifying k-mers unique to controlled viruses, maximizing specificity in pathogen detection. Analysis of selected controlled viruses revealed that most possess both a high proportion of unique 30-mers and a substantial absolute number (>10,000) of defendable 30-mers. SARS and 1918 influenza virus are notable exceptions, showing lower uniqueness due to their high sequence similarity with SARS-CoV-2 and other influenza A strains, respectively. For these cases with limited unique sequences, the database may be augmented with additional k-mers shared with selected unregulated relatives, which increases detection sensitivity while accepting a reduction in specificity. Data points for Select Agents are highlighted in red.

**Extended Data Fig. 2.**
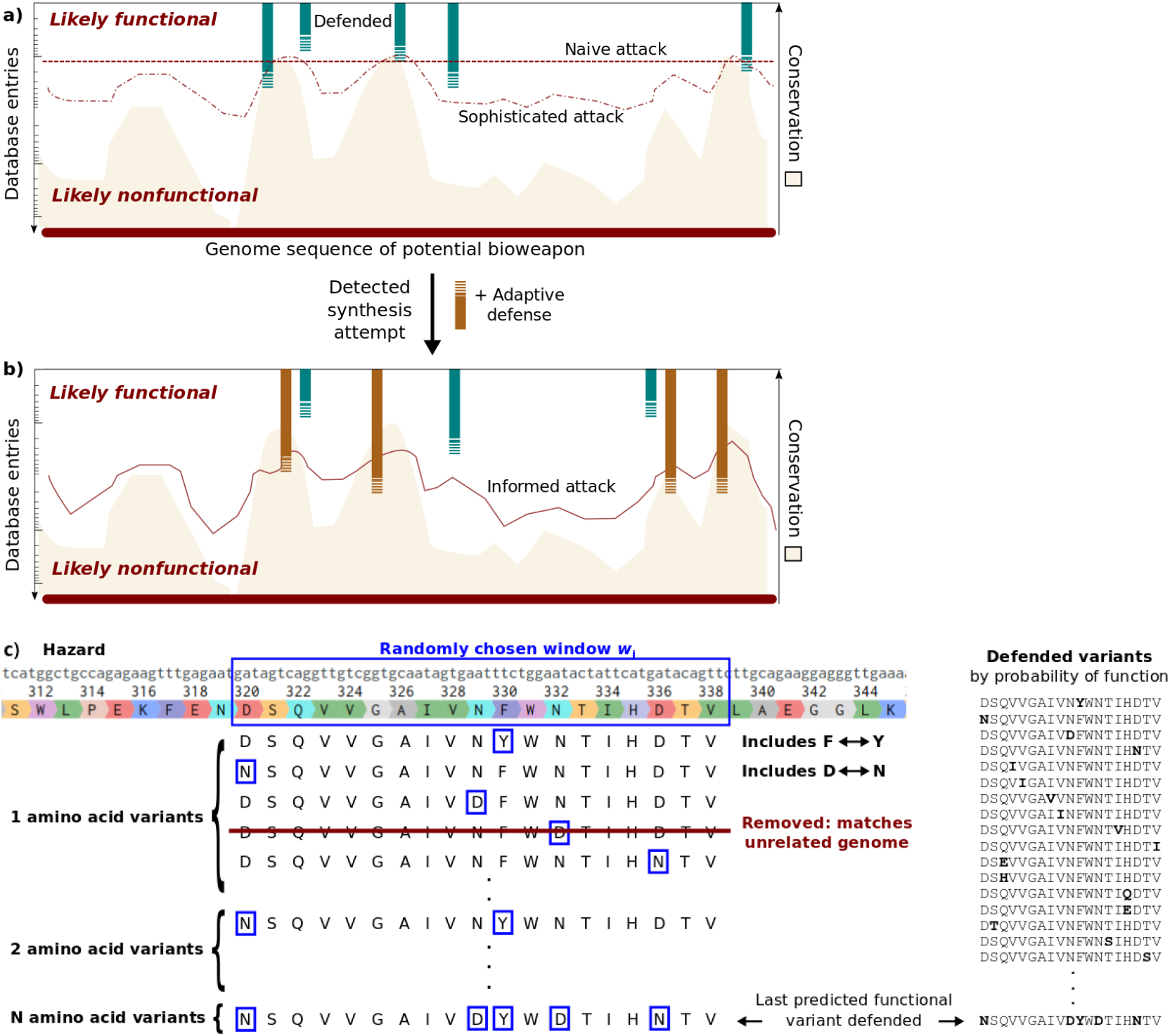
Potential attacks seeking to evade exact-match predictive screening and the possibility of adaptive defense. a) Variants from five windows in a region are included in the controlled sequence database, mostly from relatively conserved regions. Most of the variants predicted to be highly functional by a variety of different algorithms at each window are included in the database, so a naive attacker who simply introduces a moderate number of mutations at a constant rate is both highly likely to be detected and risks generating a nonfunctional hazard due to the accumulated fitness cost. A sophisticated attacker may try to tune the number of mutations to the likelihood of obtaining a functional sequence across regions, thereby maximizing the chance of evading screening while preserving function. Their chances improve with the superiority of their variant prediction capabilities relative to the defender, but they still must trade off the risk of generating a nonfunctional hazard against being randomly detected upon picking a database variant at one of the five protected windows. b) If multiple attacks on a particular hazard are detected, the system can adaptively add new windows and defend more variants at each window, precluding informed attacks based on probing or database interrogation. Windows may also be rotated in and out. c) A controlled sequence database might include predicted functional variants of quasi-randomly chosen windows comprising 20-amino-acid peptides from hazardous proteins or 30- or 42-base-pair DNA/RNA sequences from the noncoding regions of hazardous genomes. Variants matching unregulated sequences in GenBank are removed. The random adversarial threshold—the probability that an order placed by an attacker with perfect knowledge of the fitness landscape but ignorant of which sequences are defended will be detected—increases as variants are added to the database. The exact method used to predict the function of variants in order to generate the list can be randomized across several prediction methods to prevent adversaries from predicting the contents of the controlled sequence database.

**Extended Data Fig. 3.**
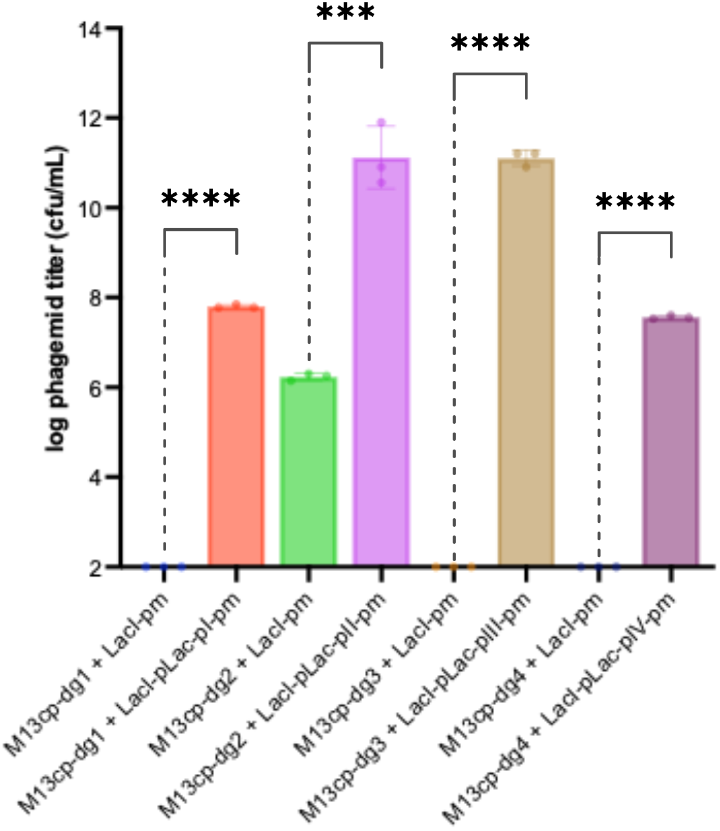
Fitness costs of loss-of-function mutations in M13 genes bearing windows. The fitness of phagemids encoding filamentous phage genes I-IV (LacI-pLac-pI-pm, LacI-pLac-pII-pm, LacI-pLac-pIII-pm, and LacI-pLac-pIV-pm, respectively) was quantified in cells carrying helper plasmids deleted for the gene in question (M13cp-dg1, M13cp-dg2, M13cp-dg3, or M13cp-dg4). In all cases, cells extruded phagemid particles when induced with 1 mM IPTG, as measured by infection of recipient cells with 3 independent biological replicates. Data from helper cells transformed with phagemids lacking the wild-type phage genes are provided for comparison. Phagemid titers below the limit of detection (100 cfu/mL) are plotted at the limit of detection. For all genes, loss of function reduced fitness by over 100-fold, which is greater than the reduction to fitness 0.05 of wild-type designated as too costly for the most contagious human virus to spread. The difference in maximum titers suggests that the optimal level of each protein differs from the level produced upon induction, which may affect the relative fitness of variants. Notably, phagemid overproduction may artificially increase the measured fitness of variants with reduced activity when excess activity is costly, resulting in an apparent mutant fitness greater than wild-type. Data plotted as mean ± standard deviation.

**Extended Data Fig. 4.**
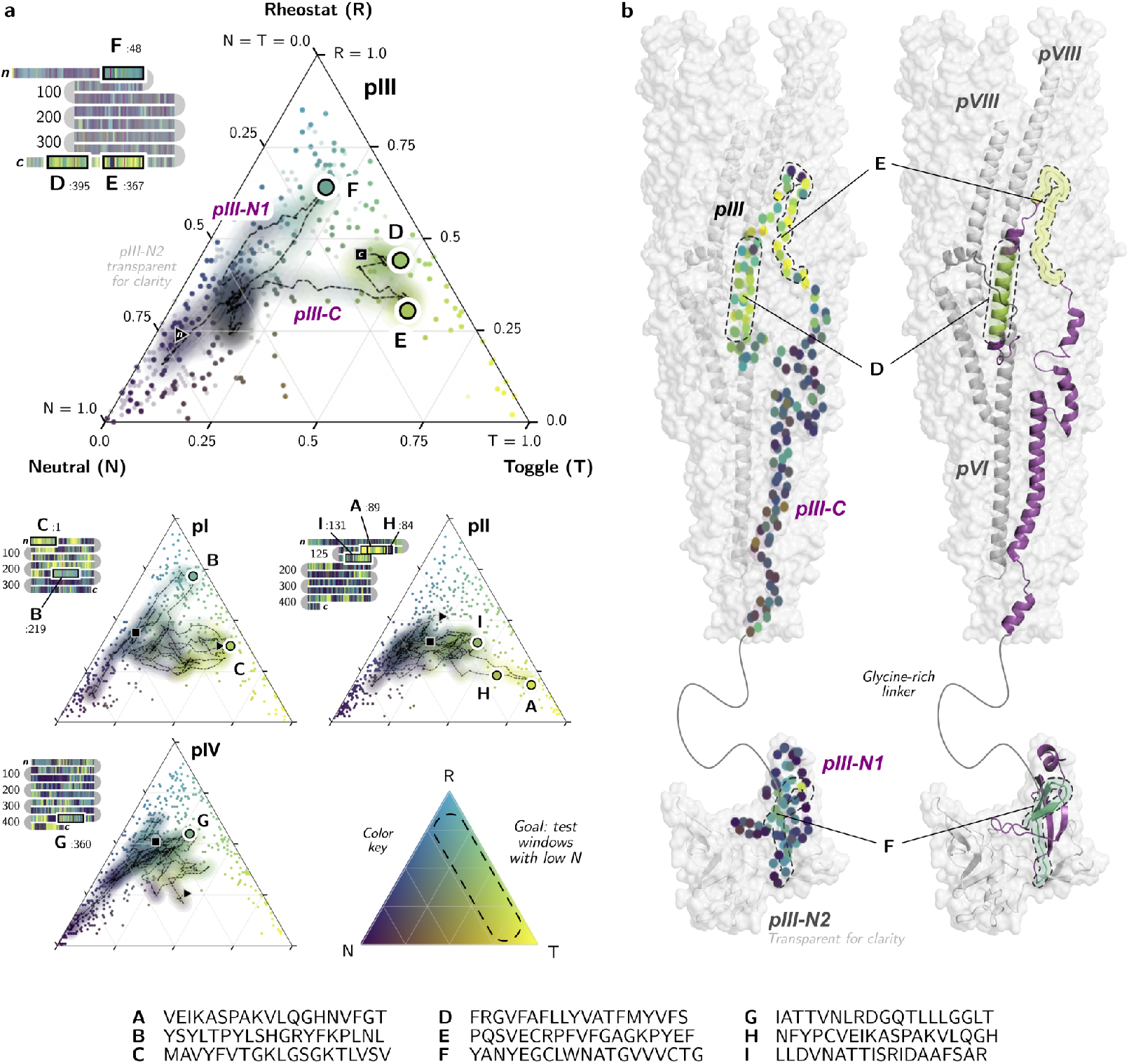
fuNTRp window analyses for M13 bacteriophage proteins. **a)** Ternary plots and color-bar snake plots show the probabilities that each residue of M13 phage proteins pI-pIV are neutral (N, purple), toggle (T, yellow), or rheostat (R, cyan), N + T + R = 1 (color key at bottom). The nine defended windows are highlighted (A-I, legend at bottom). Ternary plot and snake plot of protein pIII enlarged to show detail. On ternary plots, scatter points show NTR scores for all residues in each protein, while 19-residue moving average (dotted trace) from n-terminus (triangle) to c-terminus (square) shows local average NTRs. 19-residue windows were optimized on the basis of average NTR scores, meaning that all possible window choices fall on the dotted trace. Windows chosen for testing minimized neutral scores (N≈0, right diagonal edge), but varied in proportion of toggle *vs*. neutral. Colors of dots for defended windows represent their average NTRs. **b)** 3D structure of pIII in context of assembled phage virion tip (surfaces)^52,53^. Dots colored by NTR (left) indicate alpha carbon atoms. Ribbon representation (right) shows the structure of pIII (purple), with defended windows in pIII (D-F) highlighted, colored by average NTR. Using a low neutral score as a proxy for functional importance, minimum-neutral windows appear to correspond to regions with many contacts with nearby proteins, pVI (gray) and pVIII (light gray) in this case (D, E), and binding sites (F).

**Extended Data Table 1.**
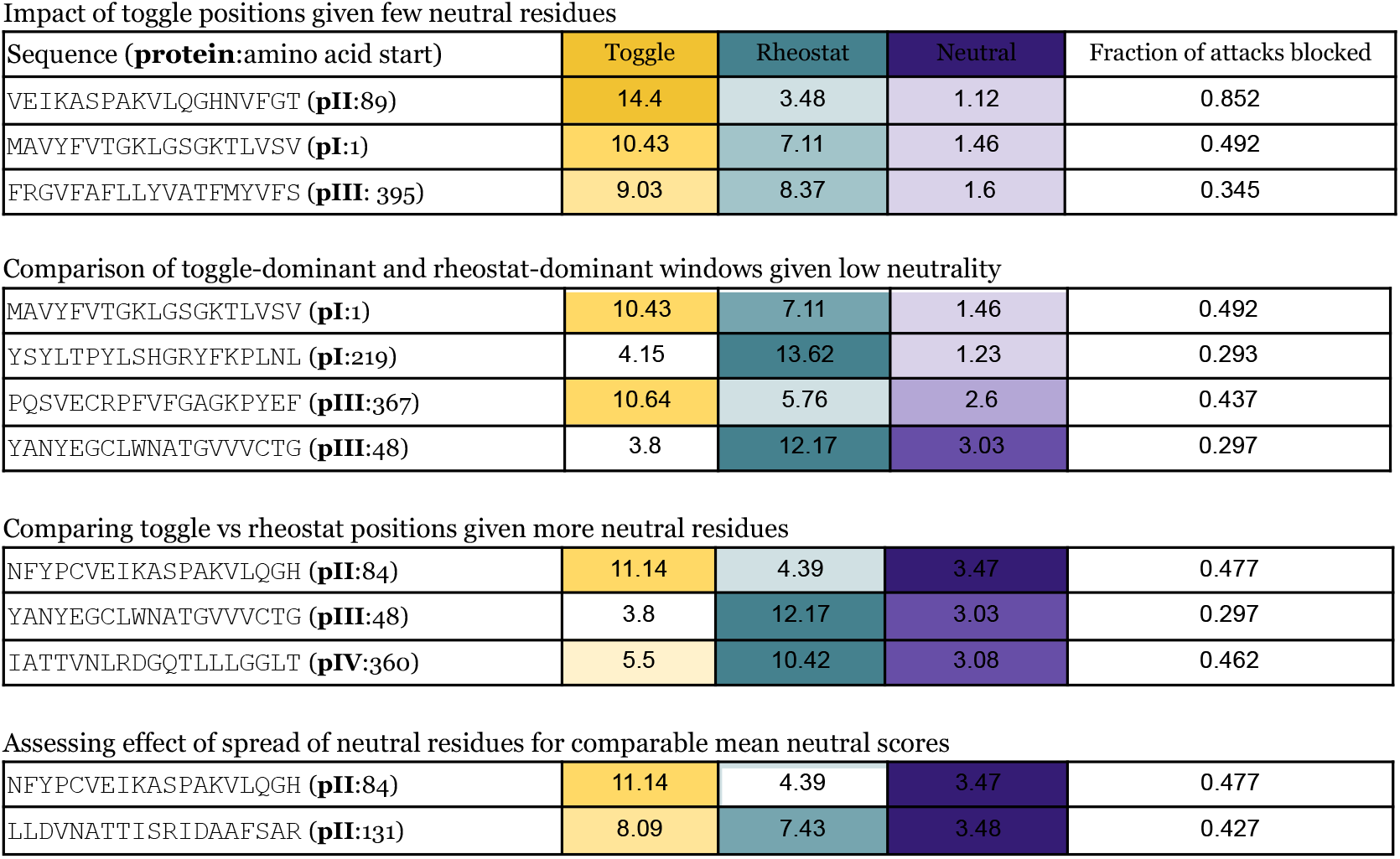
Strategic selection of windows to assess impact of funNTRp attributes on defensibility. Windows from M13 phage proteins were chosen to compare how different fuNTRp scores affect the fraction of combinatorial attacks blocked when screening 10^6^ predicted variants at each window. Colors highlight values of interest. The top section shows that with few neutral residues, more toggles (purple) increase defensibility over more rheostats (cyan). The middle section compares toggle-dominant to rheostat-dominant windows. The bottom section compares windows with comparable mean neutral scores that are either spread out or concentrated. Some windows are included multiple times to enable comparisons. This strategic selection of windows provided insights into optimizing the use of funNTRp outputs for identifying highly defensible sequences when generalizing this approach to populate the controlled sequence database used by the full exact-match predictive system for screening DNA synthesis orders. Note that some windows are repeated for convenience, and two of the windows partly overlap.

**Extended Data Fig. 5.**
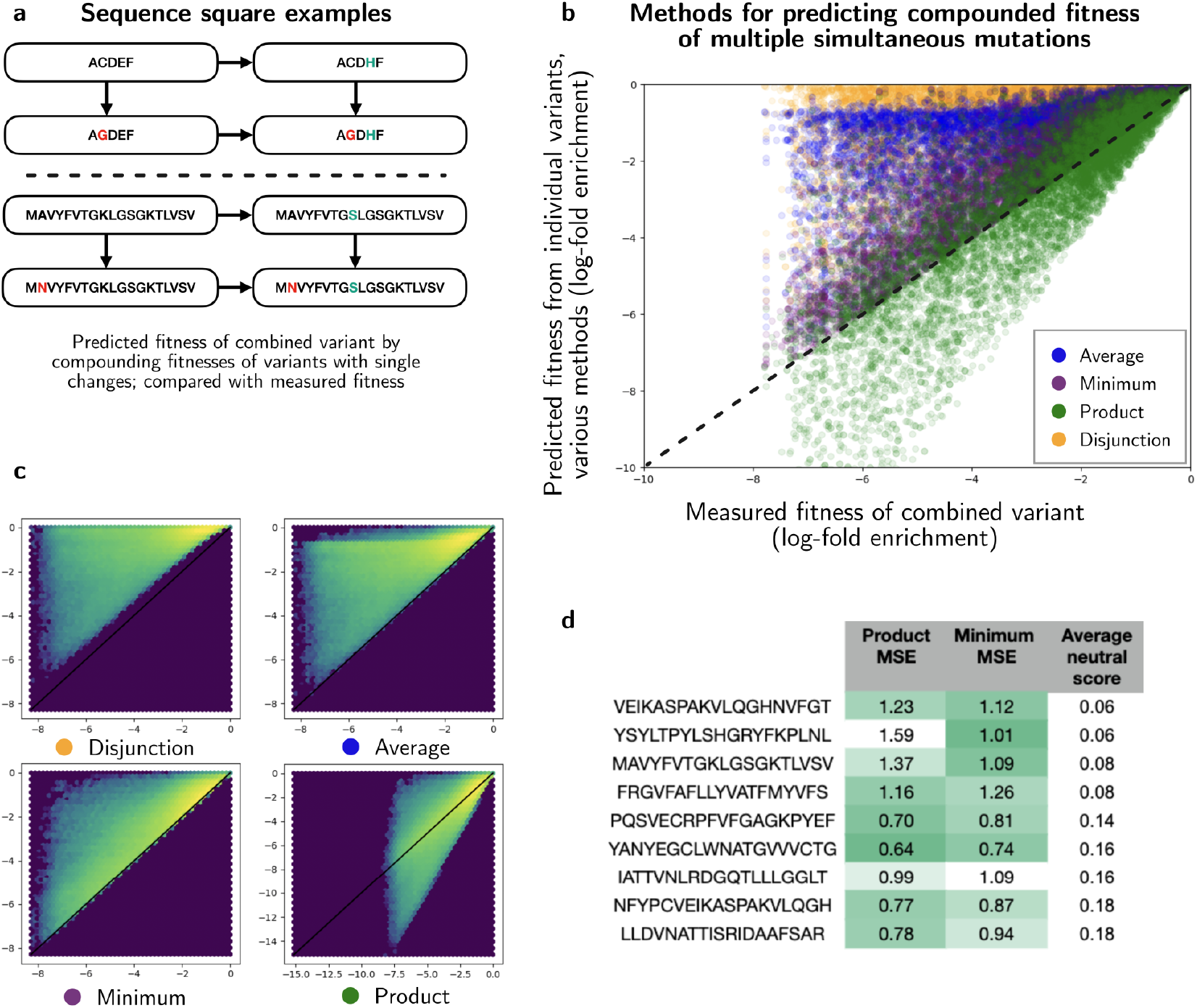
Effects of combinatorial fitness reductions. For its security claims, our analysis of exact-match predictive search assumes that fitness reductions due to individual mutations will tend to compound when such mutations are present simultaneously. Fitness of combined mutations are estimated by various methods. a) Sequence “squares” are shown, representing a variant, its orthogonal single mutants, and their combined double mutant. Top: A conceptual instance. Bottom: A sequence square from the dataset with variants of the fragment MAVYFVTGKLGSGKTLVSV. b) Fitness models for the fragment VEIKASPAKVLQGHNVFGT are assessed, comparing the Average, Minimum, Product, and Disjunction (f=1-(1-f_1_)(1-f_2_)) of 1,000 single-mutant variants against their double-mutants’ fitnesses. Dotted line denotes perfect correlation. Heuristically, Product models statistically independent changes and Minimum models breaking changes. c) Hex heat map of the fitness methods for 10^6^ sequence squares from the fragment VEIKASPAKVLQGHNVFGT shows highest concentration near the diagonal for the Minimum and Product methods. Across all fragments, Minimum and Product consistently outperformed others by mean square error (MSE). d) Fragments sorted by neutral score. For fragments with low neutral scores, expected to be regions where breaking changes are likely, the Minimum method is superior by MSE. Conversely, the Product method is superior for fragments with high neutral scores, suggesting that small independent fitness impacts can be multiplicatively combined to estimate the total fitness reduction.

**Extended Data Fig. 6.**
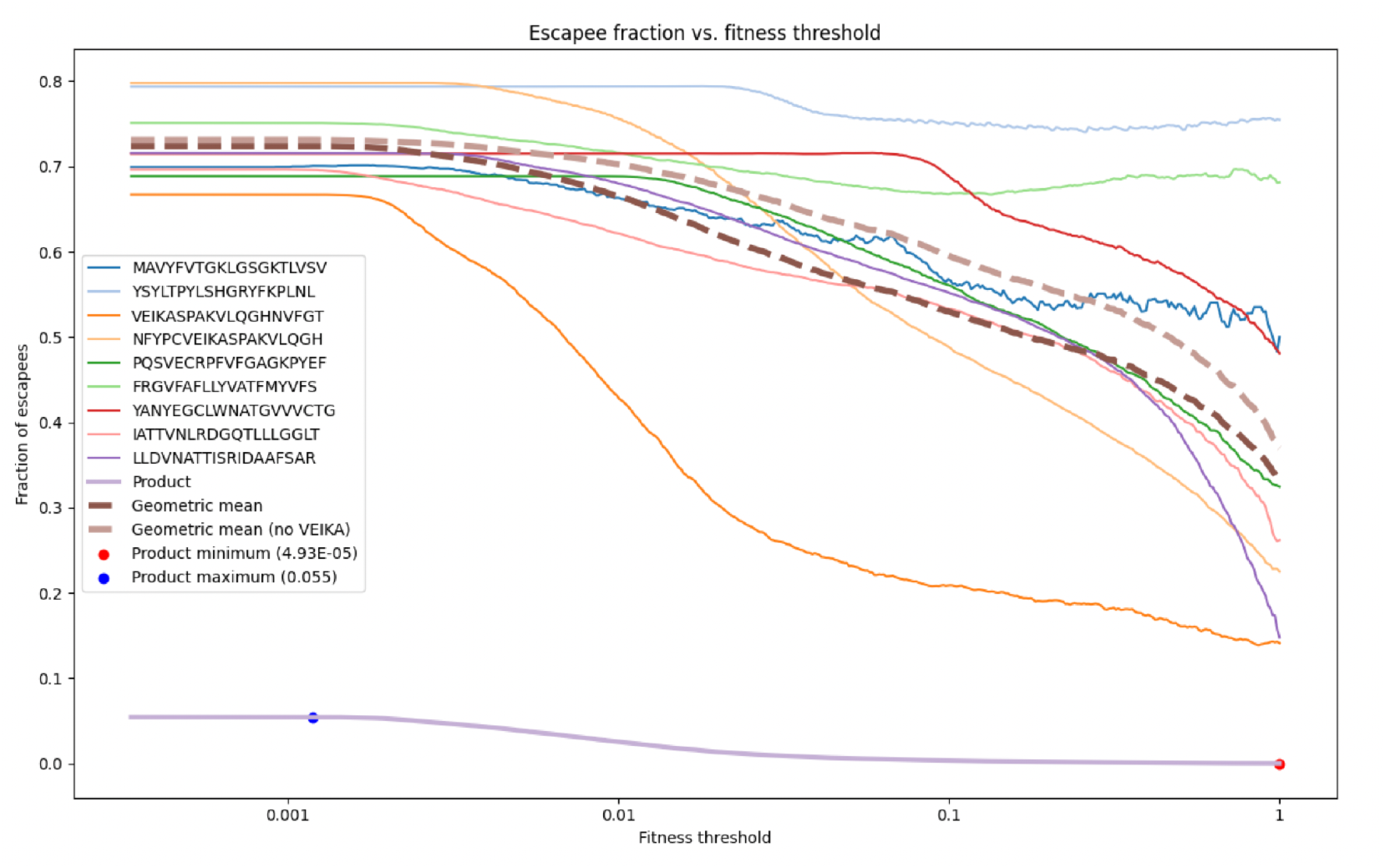
Fraction of undetected attacks against minimum fitness thresholds. The adversary is assumed to possess perfect knowledge of variant fitness out to six amino acid mutations relative to the 19-amino-acid wild-type subsequence. The horizontal axis depicts the tolerance level of the attacker to fitness hits: more damaging hits and permissive fitness cutoffs are towards the left, while higher-performing but correspondingly restrictive fitness cutoffs are on the right. Vertical axis is the fraction of escapee variants that had fitness higher than the cutoff that were not in the 50,000-variant database. The overall left-to-right decreasing trend indicates that an attacker attempting to synthesize an agent with high fitness is more likely to match a functional variant in the controlled sequence database: that is, the fuNTRp+BLOSUM62 classifier accurately predicts variant fitness at the window. Some sequence windows (VEIKA) offer robust protection even at low fitness cutoffs, whereas others (YSYLT, FRGVF) protect a roughly fixed fraction of variants regardless of the fitness threshold, indicating that our classifier has limited power to distinguish more fit variants at these windows. Classifiers based on other variant effect predictors may differ in their predictive ability across distinct windows. When all nine of these windows are combined multiplicatively (bold purple), simulating a defense based on all of them, adversaries rarely predict functional variant genomes at any fitness threshold. The geometric mean (dark brown, dotted) represents the average window such that the effect of combining more than 9 windows may be extrapolated multiplicatively. The geometric mean excluding VEIKA, which may be an outlier, is also shown (light brown, dotted).

**Extended Data Fig. 7.**
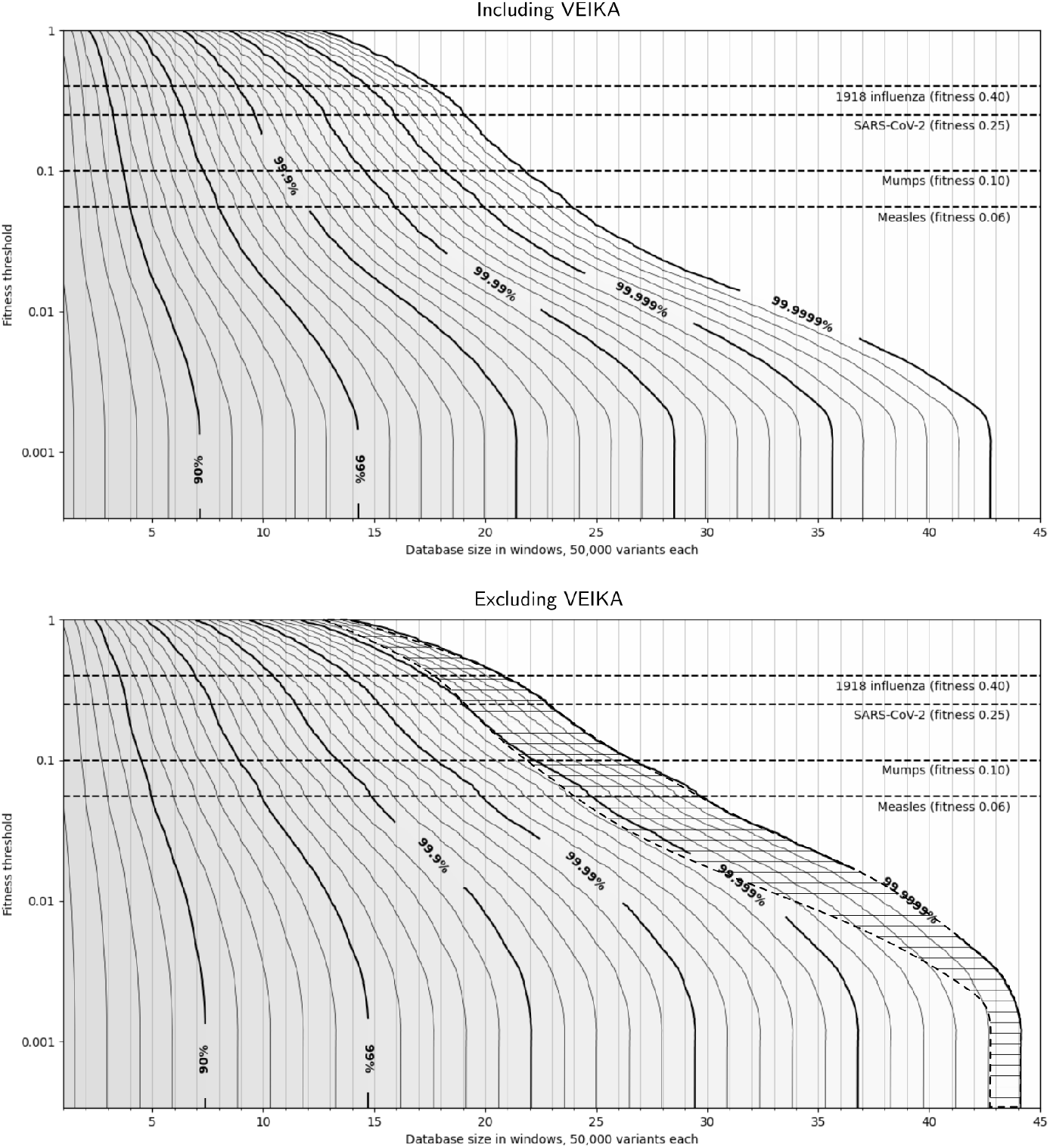
Effect of excluding outlier window. Comparison of analyses including VEIKA (top) and excluding VEIKA (bottom), which may be an outlier. Top, same data as Figure 5: original extrapolation of levels of protection from exact-match predictive search, by powers of the geometric mean of all 9 escapee curves (Extended Data Fig. 6, dark brown dotted line). The number of simulated windows, each protected with 50,000 variants, is plotted on the horizontal axis. The fitness at which the hazard is no longer functional is the vertical axis. Contours show lines of constant Random Adversarial Threshold *R*, or equal protection, as a trade-off between choices of minimum fitness tolerated and number of windows protected. Bottom: identical plot where the effect of simulated windows is extrapolated using the geometric mean of 8 escapee curves, excluding VEIKA (Extended Data Fig. 6, light brown dotted line). Shaded region shows displacement of 99.9999% contour, which moved the most. Greatest displacement was for measles. Excluding VEIKA from the estimate for the mean window, a virus as infectious as measles may require up to 6 extra windows to achieve the same *R*. Even if VEIKA is an outlier, 100 or more windows should suffice to block combinatorial attacks with high confidence.

**Extended Data Fig. 8.**
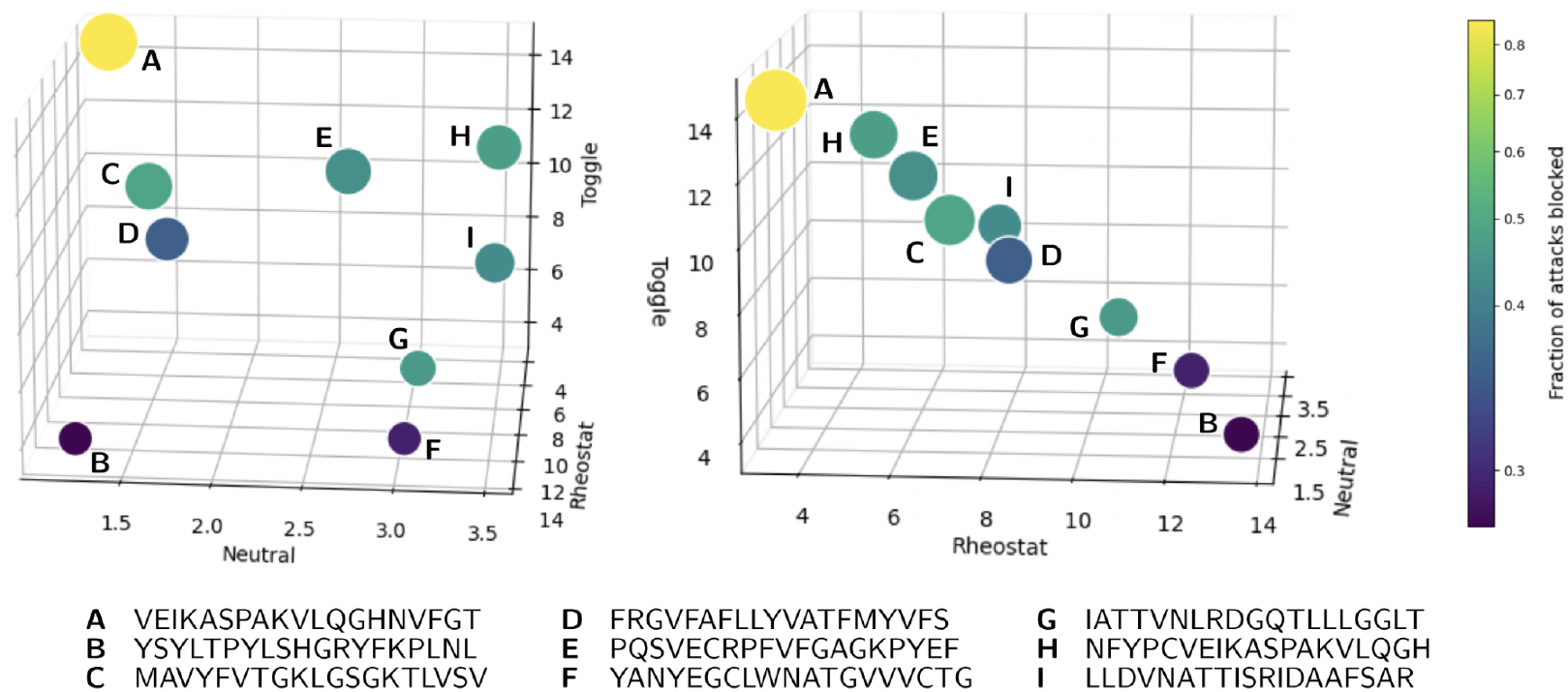
Predicted residue mutability correlates with defensibility. Points represent 19-codon windows from the M13 phage genome, plotted in 3D feature space by their average fuNTRp neutral, toggle, and rheostat scores per residue. Lighter color indicates a higher fraction of 21,000 combinatorial attacks blocked by screening 10^6^ predicted functional variants of that window. The fraction of attacks blocked serves as an experimental measurement of the window’s robustness to mutations. Windows with higher toggle scores generally blocked a higher fraction of attacks, as toggle residues are predicted to be less tolerant of mutations. funNTRp classifies residues as neutral (tolerating most mutations), toggle (highly sensitive), or rheostat (intermediate mutability). The summed scores per residue equal a constant, confining points to a plane. This analysis informed subsequent window selection to maximize screening efficacy.

**Extended Data Fig. 9.**
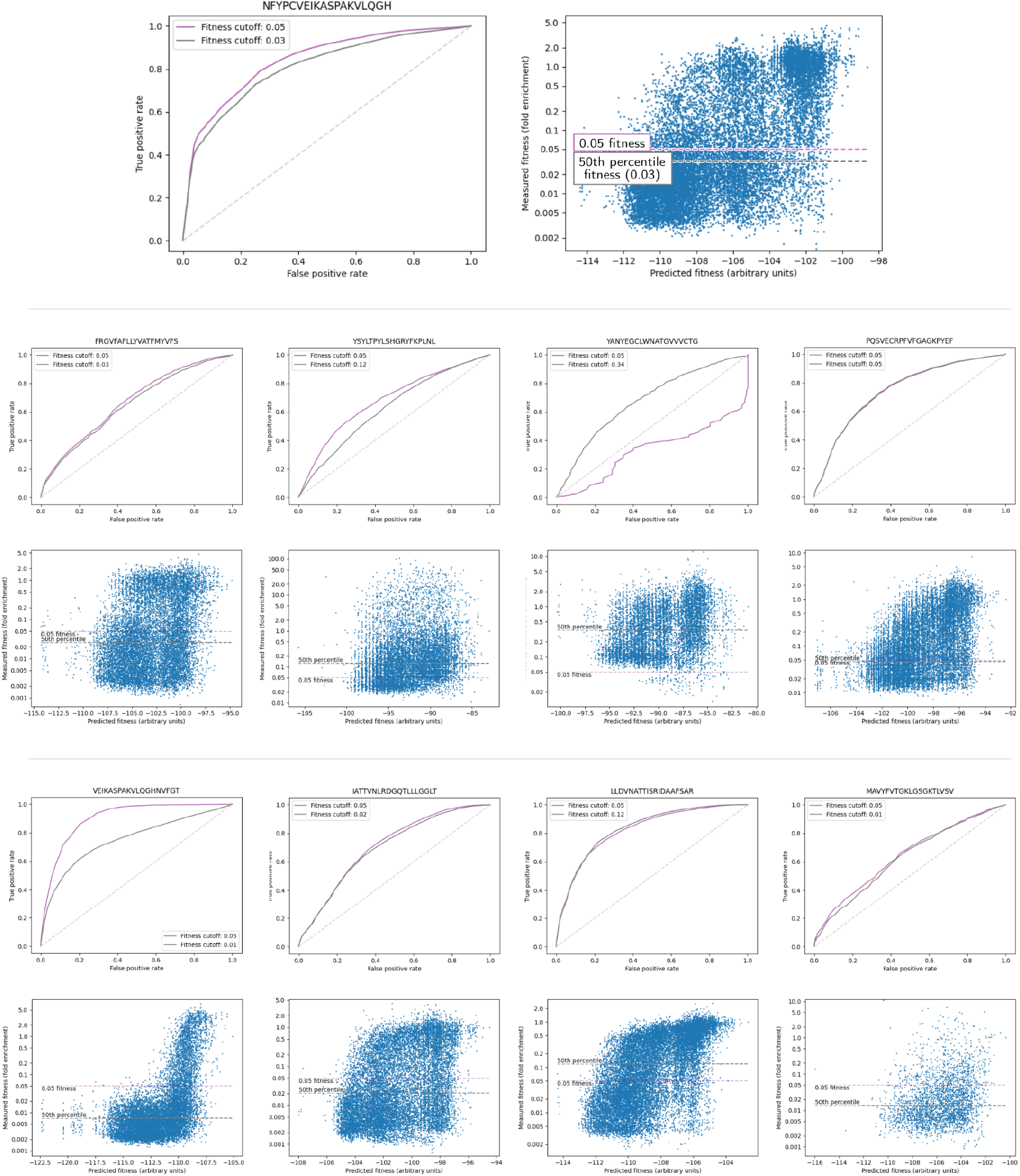
Receiver-operating-characteristic curves. Left: Receiver-operating-characteristic (ROC) curves for fuNTRp+BLOSUM62 prediction for nine 19-amino acid windows across the genome of M13 bacteriophage. ROC assesses performance of computational prediction of variant functionality for design of the controlled sequence database. Curves show true positive rate vs false positive rate in the range 0 to 1 across a range of classification thresholds. ROC curves higher toward (0, 1) indicate better performance. Dotted line from (0, 0) to (1, 1) corresponds to random guessing. Metric combines fuNTRp scores and BLOSUM62 substitution scores to predict fitness effect of amino acid substitutions. Functional variants defined as empirical fitness >0.06 wild-type (purple). ROC for median empirical fitness shown in gray. Right: fuNTRp+BLOSUM62 prediction metric vs measured relative fitness. Bottom: similar curves and point clouds for all fragments. Best-performing classifier VEIKASPAKVLQGHNVFGT demonstrates high discrimination ability, capturing >90% of functional variants at <30% false positive rates, preventing database overfill while retaining impactful variants. Poorer-performing MAVYFVTGKLGSGKTLVSV classifier still contributes protective value by correctly predicting some functional variants, while trading off less favorably with database size. YANYEGCLWNATGVVVCTG produced no meaningful ROC curve at 0.05 fitness threshold due to data sparsity; median curve indicates roughly average metric performance where data exists.

## Supplementary Information

Supplementary Figure 1 | Analyses of red-team attacks against defended windows of M13 phage.

Supplementary Table 1 | Results of unique 30-mer identification for a representative selection of controlled virus genomes in GenBank.

*Appendix A: Theoretical specificity analysis*

*Appendix B: Future-Proofing Screening Against Advances in Functional Prediction*

*Appendix C: Screening for emerging hazards*

**Supplementary Figure 1.**
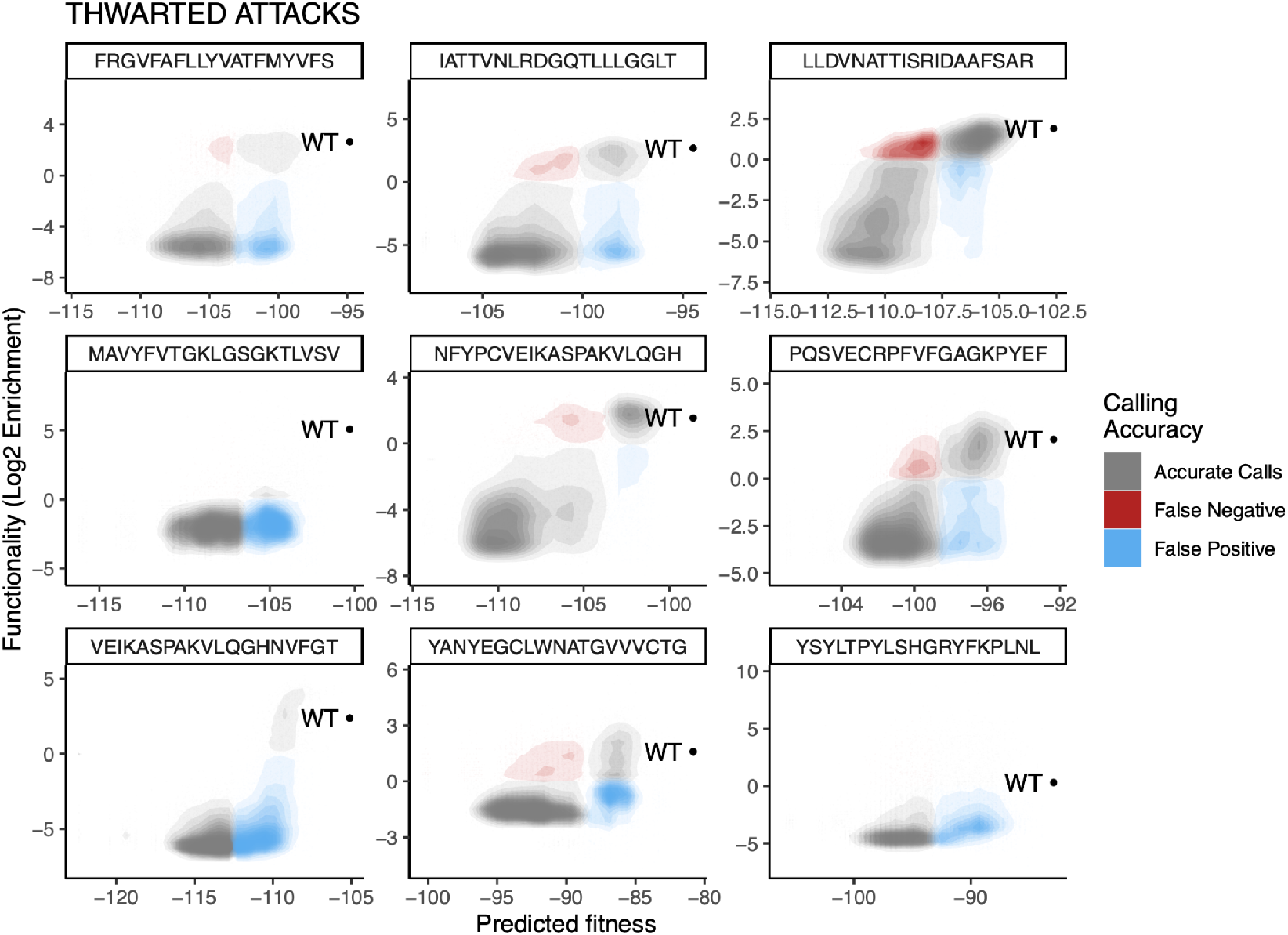
Analyses of red-team attacks against defended windows of M13 phage. Predicted fitness using fuNTRp and BLOSUM62 to estimate fitness of sequence variants is shown on the horizontal axis. Vertical axis shows variant fitness, in log fold enrichment after one round of phage replication and selection. Heat map shows density of points in each of three categories: true positives and true true negatives, grouped as “accurate calls;” false positives, which represent an instance when the predictor rated a variant’s fitness too highly and incorrectly screened it, representing an inefficiency; and false negatives, which represent successful attacks that were not thwarted at the window shown. YSYLT had negligible true positives but also no false negatives. VEIKA had almost exclusively accurate calls and false positives, representing excellent performance. LLDVNA was the worst performer by this metric in that it introduced the greatest density of false negatives; to restore security, other windows with better performance must be included in combination with this window. As this cannot be known in advance, this result highlights the importance of including many windows, which assist in driving *R* to 1.0.

**Supplementary Table 1.** Results of unique 30-mer identification for a representative selection of controlled virus genomes in GenBank.

| **Controlled virus** | **Select Agent** | **Accession number(s)** | **Total 30-mers** | **30-mers unique to virus** | **% unique 30-mers** |
| --- | --- | --- | --- | --- | --- |
| 1918 influenza | Yes | Not applicable | 13089 | 2400 | 18.3% |
| Australian bat lyssavirus | No | NC_003243.1 | 11793 | 11793 | 100.0% |
| Bokeloh bat lyssavirus | No | NC_025251.1 | 11871 | 11864 | 99.9% |
| Chapare virus | Yes | NC_010563.1,<br>NC_010562.1 | 10406 | 10399 | 99.9% |
| Chikungunya virus | No | NC_004162.2 | 11797 | 11496 | 97.4% |
| Crimean-Congo hemorrhagic fever virus | Yes | NC_005301.3,<br>NC_005300.2,<br>NC_005302.1 | 19059 | 18876 | 99.0% |
| Duvenhage virus | No | NC_020810.1 | 11977 | 11977 | 100.0% |
| Eastern equine encephalitis virus | Yes | NC_003899.1 | 11646 | 11521 | 98.9% |
| European bat lyssavirus I | No | NC_009527.1 | 11937 | 11937 | 100.0% |
| European bat lyssavirus II | No | NC_009528.2 | 11931 | 11928 | 100.0% |
| Guanarito virus | Yes | NC_005082.1,<br>NC_005077.1, | 10366 | 9756 | 94.1% |
| Hantaan virus | No | NC_005222.1,<br>NC_005219.1,<br>NC_005218.1 | 11758 | 10869 | 92.4% |
| Hendra virus | Yes | NC_001906.3 | 18205 | 18190 | 99.9% |
| Japanese encephalitis virus | No | NC_001437.1 | 10947 | 10041 | 91.7% |
| Junin virus | Yes | NC_005080.1,<br>NC_005081.1 | 10467 | 9222 | 88.1% |
| Kyasanur Forest disease virus | Yes | NC_039218.1 | 10347 | 10058 | 97.2% |
| Lassa virus | Yes | NC_004297.1,<br>NC_004296.1 | 10623 | 10592 | 99.7% |
| Lujo virus | Yes | NC_012777.1,<br>NC_012776.1 | 10294 | 10293 | 100.0% |
| Lyssavirus aravan (Avaran virus) | No | NC_020808.1 | 11889 | 11886 | 100.0% |
| West Caucasian bat lyssavirus | No | NC_025377.1 | 12249 | 12249 | 100.0% |
| Gannoruwa bat lyssavirus | No | NC_031988.1 | 11890 | 11887 | 100.0% |
| Ikoma lyssavirus | No | NC_018629.1 | 11873 | 11873 | 100.0% |
| Irkut lyssavirus | No | NC_020809.1 | 11951 | 11947 | 100.0% |
| Lleida bat lyssavirus | No | NC_031955.1 | 11902 | 11901 | 100.0% |
| Mokola lyssavirus | No | NC_006429.1 | 11911 | 11898 | 99.9% |
| Lyssavirus Ozernoe | No | NC_025408.1 | 11951 | 11951 | 100.0% |
| Shimoni bat lyssavirus | No | NC_025365.1 | 12016 | 12013 | 100.0% |
| Taiwan bat lyssavirus | No | NC_055474.1 | 11959 | 11956 | 100.0% |
| Machupo virus | Yes | NC_005079.1,<br>NC_005078.1 | 10577 | 10577 | 100.0% |
| Marburgvirus | Yes | NC_001608.3 | 19082 | 19081 | 100.0% |
| Middle East respiratory<br>syndrome coronavirus<br>(MERS-CoV) | No | NC_019843.3 | 30090 | 27599 | 91.7% |
| Mpox virus | Yes | NC_063383.1 | 196811 | 20035 | 10.2% |
| Murray Valley encephalitis<br>virus | No | NC_000943.1 | 10985 | 10866 | 98.9% |
| Nipah virus | Yes | NC_002728.1 | 18217 | 18091 | 99.3% |
| Omsk hemorrhagic fever virus | Yes | NC_005062.1 | 10758 | 10596 | 98.5% |
| Oropouche virus | No | NC_005776.1,<br>NC_005775.1,<br>NC_005777.1 | 11898 | 11677 | 98.1% |
| Rabies -virus | No | NC_001542.1 | 11903 | 11889 | 99.9% |
| Rift Valley fever virus | Yes | NC_014397.1,<br>NC_014396.1,<br>NC_014395.1 | 11892 | 11847 | 99.6% |
| Sabiá virus | Yes | NC_006313.1 | 10441 | 10430 | 99.9% |
| Saint Louis encephalitis virus | No | NC_007580.2 | 10941 | 10913 | 99.7% |
| Severe acute respiratory<br>syndrome coronavirus (SARS) | Yes | NC_004718.3 | 29722 | 5971 | 20.1% |
| Sin Nombre virus | No | NC_005217.1,<br>NC_005215.1,<br>NC_005216.1 | 12230 | 10989 | 89.9% |
| Variola virus | Yes | NC_001611.1 | 184760 | 50996 | 27.6% |
| Venezuelan equine encephalitis<br>virus | Yes | NC_001449.1 | 11415 | 10929 | 95.7% |
| Western equine encephalitis<br>virus | No | NC_003908.1 | 11455 | 11139 | 97.2% |
| West Nile virus | No | NC_009942.1 | 11000 | 10890 | 99.0% |
| Zaire ebolavirus | Yes | NC_002549.1 | 18930 | 18877 | 99.7% |
| Yellow fever virus | No | MF405338.1 | 10366 | 10333 | 99.7% |
| Andes virus | No | NC_003468.2,<br>NC_003467.2,<br>NC_003466.1 | 12017 | 10329 | 86.0% |

### Appendix A: Theoretical specificity analysis

Curation ensures that exact-match predictive search will never flag known unregulated sequences, but random matches to novel sequences will occur at a frequency determined by the total amount of novel DNA synthesized and the number of sequences in the database. The probability that any translation of a random 60-mer will match a specific 20 amino acid peptide is 8.0×10^-25^. If 10^15^ unique 60-mers of oligonucleotides will be synthesized in 2035^54^, we expect approximately one random false alarm for the entire world’s DNA synthesis in that year. Similar numbers are obtained for DNA screening. While biological sequences are far from isotropic^55^, almost all orders comprise or encode known sequences. Therefore, random false alarms will overwhelmingly occur in oligonucleotide libraries for experiments such as deep mutational scanning and directed evolution; removing a random oligo from a library of many thousands or millions is not expected to impact the results of such an experiment. Empirical measurements of specificity on datasets of real-world orders are detailed in the companion paper^40^.

The expected false-alarm (FA) rate per year depends on k-mer size and translated window size. For DNA of length k and peptides of length k/3,

FA_DNA_(k) = (novel k-mers/yr) × (k-mers in *D*) / ((1 forward + 1 backward) × 4^k^)

FA_AA_(k) = (novel DNA windows/yr) × (windows and variants in D) / [(1 forward + 1 backward) × (20 amino acids × 61 / 64 codons)^(k/3)^]

### Appendix B: Future-Proofing Screening Against Advances in Functional Prediction

The experimental results in this study rely on simple prediction tools (fuNTRp with BLOSUM62 substitution matrices) to counter sophisticated combinatorial attacks. A key concern is whether improvements in variant effect prediction could grant adversaries superior predictive capabilities. However, multiple factors ensure defenders will retain their advantage:

- **Asymmetric computational resources.** Real-world screening systems would leverage significantly greater computational power over a longer period than typical adversaries, enabling the use of more sophisticated, compute-intensive prediction models.
- **Privileged access to prediction tools.** Organizations maintaining screening databases can be expected to learn about and gain early access to state-of-the-art prediction tools and/or have greater in-house proficiency in adopting and applying new tools than adversaries, allowing defensive systems to integrate superior models faster than average after they become widely available.
- **Dynamic database updates.** Unlike the static database used in our experiments, operational screening systems can continuously incorporate new prediction models, regenerating or augmenting databases to counter expected improvements in adversarial strategies.
- **Optimized window selection.** Improved prediction tools enable defenders to strategically choose windows, balancing unpredictability with defensibility as desired, whereas adversaries will always be obligated to mutate all windows to avoid detection.
- **Combinatorial advantage.** Even with perfect prediction capabilities, adversaries must evade detection across multiple windows while maintaining function. Defenders need only identify functional variants in a small subset of windows to achieve high detection rates, maintaining an intrinsic advantage.

Our results confirm this asymmetry: simple prediction models successfully detected all functional ProteinMPNN redesigns. As predictive methods advance on both sides, this fundamental imbalance ensures that defenders remain ahead as long as they continue to apply sufficient computational resources and implement continuous updates.

### Appendix C: Screening for emerging hazards

Exact-match predictive search is compatible with cryptographic approaches capable of obscuring the identities of entries in an emerging controlled sequence database. Such a system might be warranted if nations decide that some threats should not be made credible by highlighting them in public screening databases, although it would require security validation before implementation. In principle, such a system would enable a researcher concerned about an emerging potential biological weapon to safely take action to restrict global access without creating information hazards^56,57^ by securely conveying their concern to the system operators. The operators could add sequences from the hazard to the encrypted emerging controlled sequence database without requiring further disclosure. If implemented, at least a dozen times as many genes or genomes would be chosen as “decoys”: related genes or agents that might seem to pose a threat, but are not actually of concern. The use of decoys can ensure that anyone who finds a match to the database will learn only that it corresponds to a plausible-seeming threat, not that it is a credible weapon. This would ensure that adversaries cannot learn whether a specific hazard can be used to build a weapon of mass destruction by attempting to screen it, e.g. by ordering it or synthesizing it on a benchtop. A detailed explanation of the cryptographic approaches required is available elsewhere^40^. Emerging hazards would only be added to the database after multiple years of testing to verify that the implementation is secure.

## Notes

### Summary of Updates

New Fig. 1, text updated; author affiliations updated.

